# Intrinsic disorder in the T cell receptor creates cooperativity and controls ZAP70 binding

**DOI:** 10.1101/2020.05.21.108662

**Authors:** Lara Clemens, Omer Dushek, Jun Allard

## Abstract

Many immunoreceptors have cytoplasmic domains that are intrinsically disordered (i.e., have high configurational entropy), have multiple sites of post-translational modification (e.g., tyrosine phosphorylation), and participate in nonlinear signaling pathways (e.g., exhibiting switch-like behavior). Several hypotheses to explain the origin of these nonlinearities fall under the broad hypothesis that modification at one site changes the immunoreceptor’s entropy, which in turn changes further modification dynamics. Here we use coarse-grain simulation to study three scenarios, all related to the chains that comprise the T Cell Receptor. We find that, first, if phosphorylation induces local changes in the flexibility of TCR *ζ*-chain, this naturally leads to rate enhancements and cooperativity. Second, we find that TCR CD3*ϵ* can provide a switch by modulating its residence in the plasma membrane. By constraining our model to be consistent with the previous observation that both basic residues and phosphorylation control membrane residence, we find that there is only a moderate rate enhancement of 10% between first and subsequent phosphorylation events. And third, we find that volume constraints do not limit the number of ZAP70s that can bind the TCR, but that entropic penalties lead to a 200-fold decrease in binding rate by the seventh ZAP70, potentially explaining the observation that each TCR has around six ZAP70 molecules bound following receptor triggering. In all three scenarios, our results demonstrate that phenomena that change an immunoreceptor chain’s entropy (stiffening, confinement to a membrane, and multiple simultaneous binding) can lead to nonlinearities (rate enhancement, switching, and negative cooperativity) in how the receptor participates in signaling. These polymer-entropy-driven nonlinearities may augment the nonlinearities that arise from, e.g., kinetic proofreading and cluster formation. They also suggest different design strategies for engineered receptors, e.g., whether or not to put signaling modules on one chain or multiple clustered chains.

**STATEMENT OF SIGNIFICANCE:** Many of the proteins involved in signal processing are both mechanically flexible and have multiple sites of interaction, leading to a combinatorial complexity making them challenging to study. One example is the T Cell Receptor, a key player in immunological decision making. It consists of 6 flexible chains with 20 interaction sites, and exhibits nonlinear responses to signal inputs, although the mechanisms are elusive. By using polymer physics to simulate the T Cell Receptor’s chains, this work demonstrates that several of the nonlinear responses observed experimentally emerge naturally due to constraints on the chains that change their entropy. This work points to new avenues to modulate signaling proteins for therapeutics by modulating their mechanical flexibility and spatial extent.

## INTRODUCTION

Challenging the tenet that “structure determines function”, more than 40% of human proteins contain intrinsically disordered regions longer than 30 amino acids (1, 2). Intrinsically disordered regions appear as linkers between globular domains (3), associated with the cytoskeleton (4, 5), or in signaling networks (6, 7). These regions often include sites for binding and post-translational modification, suggestions they serve a purpose beyond as passive tethers (3, 8, 9). Additionally, the length of the domains themselves influences their interactions, affecting binding kinetics and catalytic performance (10–13).

One example is offered by the T Cell Receptor (TCR) (14–16). TCR has eight subunits, six of which contain intrinsically disordered cytoplasmic tails: ζ (2 per TCR), ϵ (2 per TCR), δ, and γ. The tyrosines on these tails are organized in pairs called ITAMs (immunoreceptor tyrosine-based activation motifs) and become phosphorylated by a kinase (LCK) upon extracellular ligand binding to the TCR. Phosphorylation of the ITAMs allows a cytosolic signaling molecule, ZAP70, to bind to the phospho-tyrosines (17) and propagate the activation signal (18). Despite lacking structure in the intracellular cytoplasmic tails, the TCR itself endows the signaling pathway with what we term nonlinearities, raising specific questions:

1. The *ζ* chains exhibit cooperative phosphorylation (19, 20), which can endow systems with ultrasensitivity (21). A hypothesis for this nonlinearity is in local changes in flexibility upon phosphorylation, as shown in Fig. 1Bii and as observed for other intrinsically-disordered proteins (8, 22). Would this local structuring be sufficient to explain the cooperativity? If so, this may explain why many immune receptors have disordered cytoplasmic tails containing multiple phosphorylation sites (10, 23).
2. The *ϵ* chain is known to associate with the inner leaflet of the membrane (24, 25), as are many other immune receptor chains including *ζ* (26), CD28 (27) and BCR (28). This has been hypothesized to sequester the tyrosines to guard against phosphorylation (Fig. 1Biii) before a signal is initiated by an antigen. But, given that tyrosine phosphorylation is one of the first steps of T cell activation, how do the first tyrosines become phosphorylated in order to induce membrane dissociation?
3. Finally, although the full TCR complex has 10 ITAMs, only 6 ZAP70 molecules associate with the receptor at one time (29). What property sets the limits of simultaneous occupation (Fig. 1Biv)?

**Figure 1:**
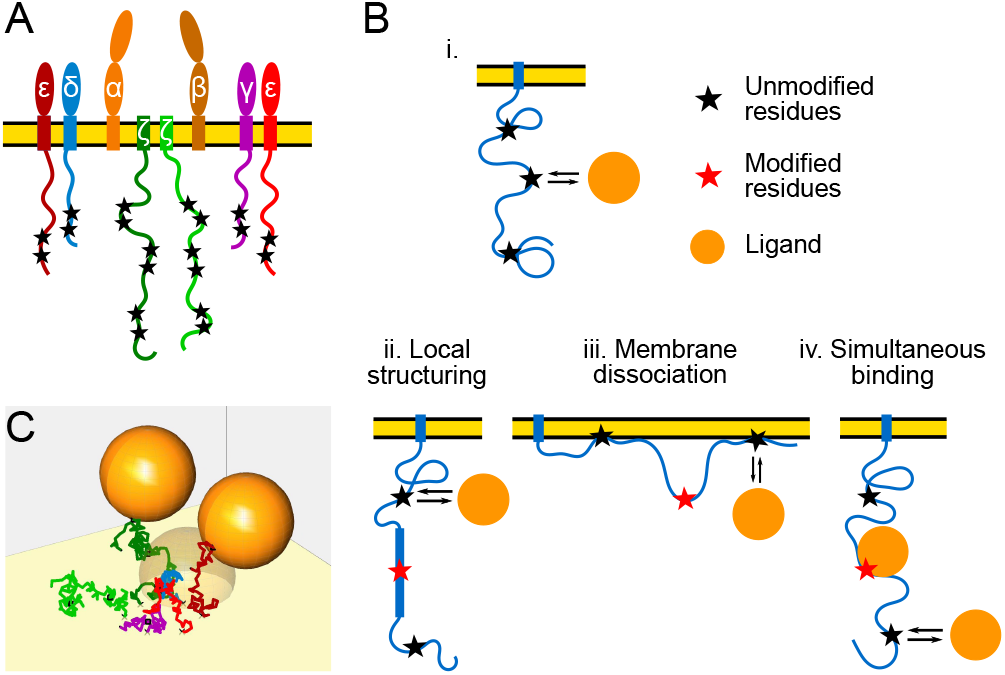
Possible consequences of membrane-bound disordered protein interacting with ligand. (A) Cartoon of T cell receptor complex. (B) (i) Interaction of ligand (orange circle) with membrane-bound disordered protein with multiple binding sites (black stars) where (ii) binding causes local stiffening near post-translational modification (e.g. phosphorylation; red stars), (iii) phosphorylation reduces membrane-association of the polymer or (iv) the ligand remains bound while more ligands attempt to bind simultaneously. (C) Snap-shot of residue-scale computational model of T cell receptor.

Computational study of the TCR is challenging, not only because it is large and membrane-bound, but the intrinsic disorder introduces a large configuration space that is explored on microseconds timescales, making atomistic molecular dynamics methods computationally expensive. An alternative modeling approach is to use coarse-grain models from polymer physics in which the disordered regions are represented as ideal chains, with each residue represented by a particle. Despite the simplicity of the approach, “residue-scale” models have proven valuable (11, 30, 31), including for formins (12, 32) and kinesins (33). Here we use coarse-grain, residue-scale models to simulate the intrinsically-disordered regions of the T Cell Receptor, exploring the consequences of post-translational modification to answer the above questions.

We find that, first, in agreement with previous theoretical calculations (19), if phosphorylation induces local changes in the flexibility of the chain, this naturally leads to rate enhancements and cooperativity. We find the rate enhancement for a single chain with properties of *ζ* is around two-fold, with Hill coefficient around 1.8. However, the ultrasensitivity seen is only strong if the kinase is large in molecular size compared to the phosphatase.

Second, we find that CD3*ϵ* can exhibit a switch-like response to phosphorylation by modulating its residence in the membrane. The observation that both tyrosine phosphorylation and basic residue deletion change membrane residency, under the assumptions of our model, sets a requirement that there is a rate enhancement of at least 10% between first and second phosphorylation events.

And third, we ask how many ZAP70 molecules can simultaneously bind the 10 ITAMs on the 6 chains of a TCR. We find that volume constraints do not limit the number (i.e. there exists a configuration where 10 can fit), but that the entropic penalty needed to bind subsequent ZAP70s leads to a 200-fold decrease in binding rate by the seventh ZAP70, potentially explaining the observation that each TCR has around six ZAP70 molecules bound following receptor triggering. We also explore how binding rates are affected if the 10 ITAMs are distributed on different numbers of chains, a question which may be relevant for artificially engineered receptors. We find that, for parameters similar to the TCR, neighbor-chain interference dominates self-chain interference, so receptors with signaling modules on the same chain would allow faster simultaneous binding.

In all three scenarios, our results demonstrate that phenomena that change an immunoreceptor chain’s entropy (stiffening, confinement to a membrane, and multiple simultaneous binding) can lead to emergent behavior with consequences for how the receptor participates in signaling. These polymer-entropy-driven nonlinearities may augment the nonlinearities that arise from, for example, kinetic proofreading and cluster formation. They also suggest different design strategies for engineered receptors, e.g., whether or not to have one chain or multiple clustered chains.

## METHODS

### Polymer model of disordered protein and globular binding partner

We represent the TCR cytoplasmic tails as ideal disordered proteins using a θ-solvent freely-jointed chain model in which each particle represents a single residue (31). The Kuhn length is δ = 0.3 nm (25, 30). The number of segments *N* is therefore the number of residues in the tail, i.e., 113, 55, 47, and 45 for *ζ,ϵ,δ,γ*, respectively.

The chains interact with LCK and ZAP70, which for generality we refer to as the ligand. This is modeled as an idealized sphere which interacts with the chain. For each ligand, we estimate the volume of the domain of interest and calculate the radius for a sphere with the specified volume. Volume estimates are made using the molecular mass of the domains and an estimated protein density of 1.41 g/cm^3^ (34). For comparison, we also estimate volumes from measurements of crystal structures of the domains (LCK - PDB 3LCK; ZAP70 - PDB 2OQ1; CD45 1YGR) in PyMol (35–37). For LCK, we represent the kinase domain as a sphere with radius 2.1 nm (7 Kuhn lengths). For ZAP70, we represent the tandem SH2 domains as a sphere with radius 2.7 nm (9 Kuhn lengths). For CD45, we represent the tandem phosphatase domains as a sphere with radius 3.4 nm (11 Kuhn lengths) For simulations of the full TCR, the relative location of the membrane-anchor for each cytoplasmic domain is estimated from (38) (PDB 6JXR).

We compute quasi-equilibrium statistics of the chain and its ligands using the Metropolis-Hastings Algorithm, a form of Monte Carlo simulation (39, 40) that generates a distribution of configurations from the canonical ensemble. At each proposed configuration, the Metropolis algorithm computes the energy of the system and accepts or rejects based on energetic preference (specifically, interactions in Eq. 5-6 below) and hard constraints (the polymer cannot pass through the membrane or ligands, and ligands cannot pass through each other). The Metropolis algorithm proposes configurations using a perturbation size that is adaptive, and increases or decreases until the acceptance rate is 0.44 (41). We repeat configuration proposals until the sequence of samples has reached a stationary distribution, which we define when the third quarter and fourth quarter of the sequence have the same distribution according to the Kolmogorov-Smirnov statistics.

Polymer simulation code and analysis routines are available at github.com/allardjun/IntrinsicDisorderTCRModel, (DOI:10.5281/zenodo.4117255).

### Calculation of ligand binding rates from occlusion statistics

The dissociation constant of a binding reaction *K*_*D*_ ≡ *k*_off_/ *k*_on_ is influenced by changes in entropy. If a binding reaction limits the configurational freedom of the binding partners, the entropy, and thus the free energy, is reduced. We can compute the change in *K*_*D*_ between two states (e.g., fully phosphorylated compared to dephosphorylated) as (22, 42):

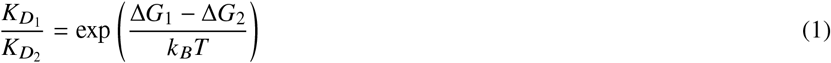

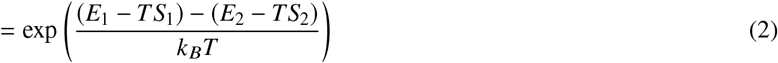

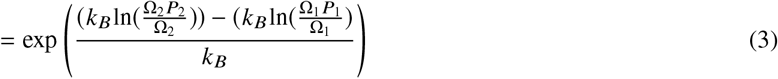

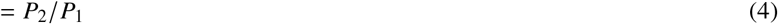

where *G* _*j*_ = *E*_*j*_ − *TS*_*j*_ is the free energy of binding in a given state, *S*_*j*_ = *k*_*B*_lnΩ_*j*_ is the entropy of binding, Ω_*j*_ is the number of microstates and *P*_*j*_ is the probability in the canonical ensemble that the configuration allows for binding. We assume Δ*E*_1_ = Δ*E*_2_, i.e., the change in energy due to ligand binding is the same, regardless of conformation. Define *P*_*occ*_ as the probability that the region of space needed by the ligand is occupied by some of the polymer, or another steric barrier. Then, 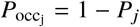, and 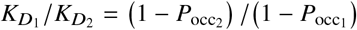 Although entropic forces could also impact unbinding of the polymer, we assume this influence to be negligible compared to the change in *k*_*on*_. Therefore, we assume that the change in *K*_*D*_ manifests as a change in *k*_*on*_. This leads to the final simple equation for attachment rates, 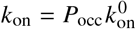, where we refer to 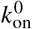 as the free-space (i.e., no occlusion) binding rate. Since we lack an estimate for 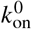 (which we expect to depend on local concentration of binding partner), we report all rates proportional to 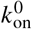

### Simulation of multi-step processes from individual rates

Given the binding rates *k*_on_, we use a Gillespie algorithm to investigate the rate-dependent behaviors of both irreversible and reversible (de)phosphorylation reactions. A matrix of occlusion probabilities, to each site in each phosphorylation state, is created from the Metropolis simulations.

For single-direction simulations, e.g., just phosphorylation, we make two calculations: 1) the probability of a specific sequence of irreversible (de)phosphorylation, and 2) the sequence-weighted average binding rate to transition between phosphostates (e.g. from one to two total phosphorylations). At the end of each run of the multi-step process, when all sites are modified, we record the event sequence and the times to transition between each step. Probabilities of each sequence are computed based on the total iterations of the algorithm. The path-weighted average binding rates are calculated as the inverse of the average transition time for a specific step. In other words, transitions between two states that are more likely are weighted higher. In the Supplemental Material, we show average binding rates that are not weighted by the probability of their path. For simulations of reversible reactions, we run until the system reaches a steady state (∼10^6^ events), and then calculate the average number of phosphorylated sites.

### Membrane interactions

Polymer-membrane interactions are modeled as potentials acting on each segment of the chain. We consider three groups of residues: tyrosines, phosphotyrosines, and basic residues. Each experiences a potential summarized in Fig. 5A. Basic residues are needed to create membrane-association (26), which we model with a piecewise parabolic potential with depth *E*_*B*0_ (43, 44),

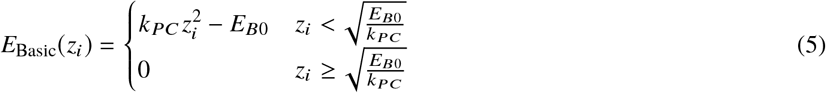

Tyrosine phosphorylation is sufficient to dissociate the polymer from the membrane (26). We therefore model phosphorylated tyrosines as having a repulsive interaction with the membrane,

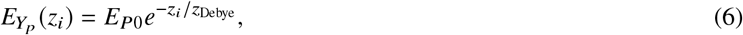

where the Debye length describes the length scale of electrostatic interactions. For cytoplasm, *z*_*Debye*_ = 1 nm and 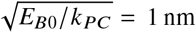 nm consistent with (31). A soft wall constraint

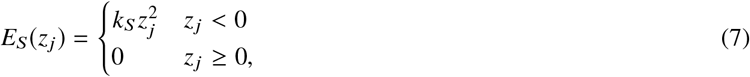

With *k*_*S*_ = 0.05*k*_*B*_*T*, is applied to the unphosphorylated tyrosines and remaining amino acids.

## RESULTS

### Site difference driven by position along chain

The residue-scale polymer model we developed simulates the T Cell Receptor as it explores its configurations, along with binding partners that we generically refer to as ligands. An example is shown in Fig. 1C and Supporting Movie SM1.

In the present section, we simulate only the *ζ*-chain. We first assume that phosphorylation does not change the polymer properties of the chain. We calculate the binding rate to each of the six tyrosines. The entropy calculations result in a relative rate from every possible reaction phosphostate, shown as the left columns in Supporting Material Fig. S1, to every phosphostate with one more phosphorylation, in the right column. From these, we obtain the phosphorylation sequences shown in Fig. 2 and Fig. 3 (pink curve, lowest in both panels).

**Figure 2:**
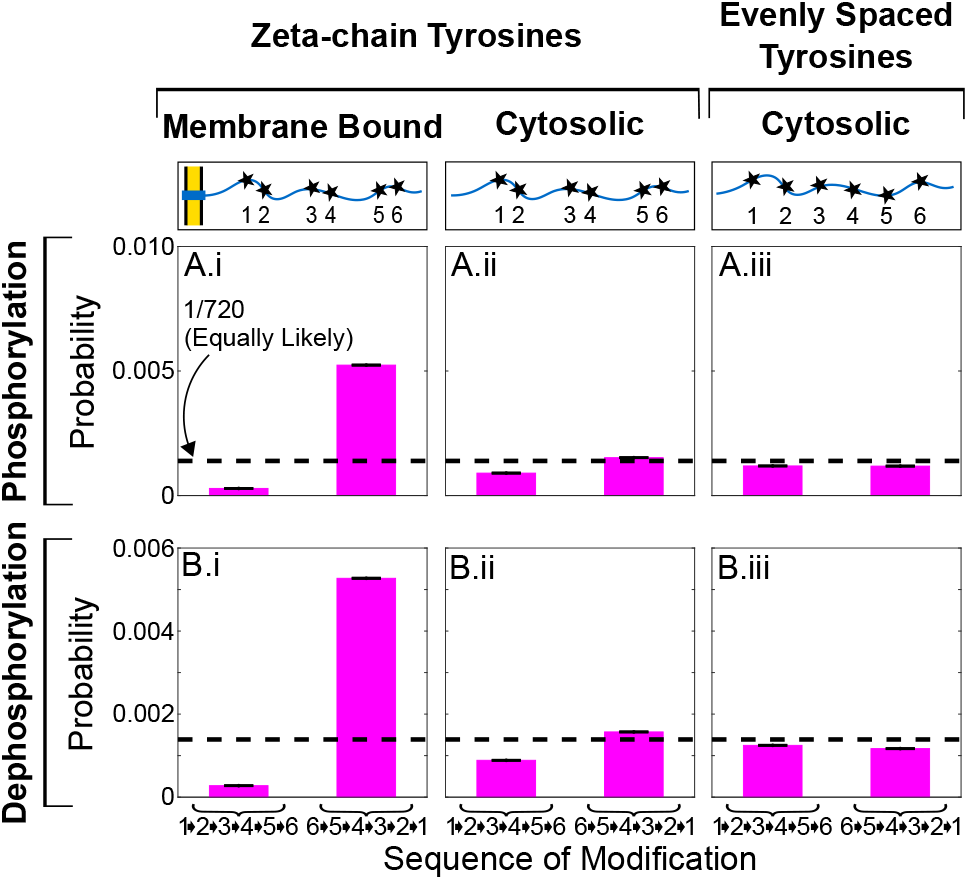
Binding rate differences imply emergent preference of (de)phosphorylation sequence. Probability of phosphorylating or (B) dephosphorylating membrane proximal-to-distal (123456) compared to membrane distal-to-proximal (654321). Each of these is shown for: (i) with a membrane, assuming sites spaced like tyrosines of *ζ*, (ii) without a membrane, assuming sites spaced like tyrosines of *ζ*, and (iii) without a membrane, assuming sites spaced evenly. Black dotted lines indicate the probability if all events were equally likely (1/6!=1/720). Error bars (A,B) represent standard error of the mean.

**Figure 3:**
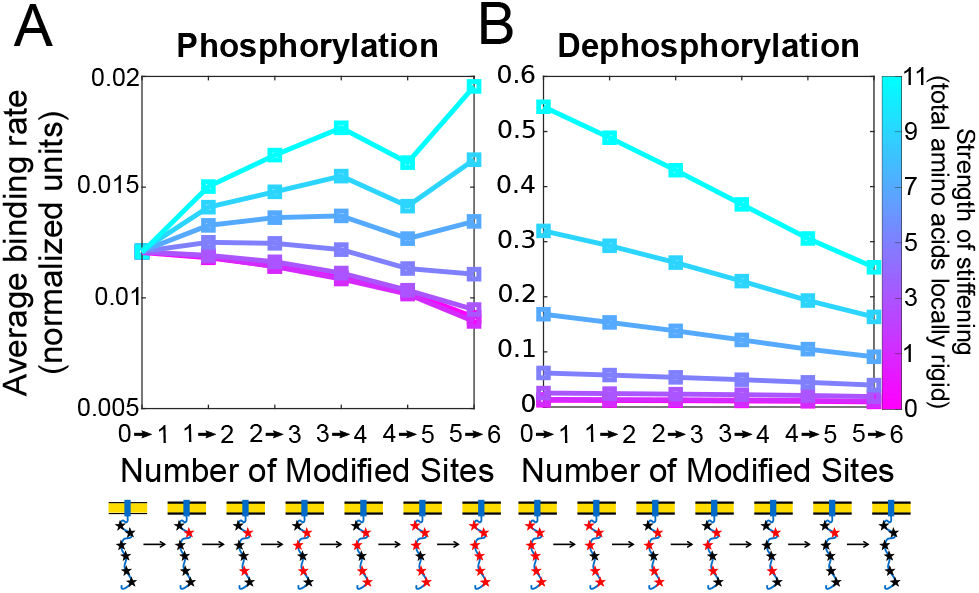
Local stiffening modulates binding rates to multi-site disordered domain. (A) Average binding rates of a kinase binding to *ζ* at different phosphorylation states for varying range of local stiffening (pink: no residues are stiffened; blue: 11 residues are stiffened, 5 on each side of site). (B) Average binding rates of a phosphatase binding to *ζ* at different dephosphorylation states for varying ranges of local un-stiffening per dephosphorylation event. For both (A) and (B), schematic below axis shows example configuration for each phosphorylation state. Kinase and phosphatase radius is 2.1nm. Rates are normalized to the free-space binding rate 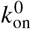.

#### Rate differences imply emergent preference of phosphorylation sequence

There are six binding sites on the *ζ*-chain, offering 720 (6 factorial) possible sequences for phosphorylation. Using a Gillespie algorithm in conjunction with the binding rates found above, we compute the probability of (de)phosphorylating in each sequence. For simplicity, Fig. 2 A.i,B.i shows only two extreme sequences: where (de)phosphorylation occurs membrane proximal-to-distal (123456, i.e., N-to-C terminus) and membrane distal-to-proximal (654321, i.e., C-to-N terminus). For membrane-bound *ζ*-chains, there is an approximately 10-fold increase in the probability of binding membrane distal-to-proximal compared to membrane proximal-to-distal for both the kinase and phosphatase. This suggested to us that disordered domains in the presence of a membrane can have an emergent preference of phosphorylation sequence.

To test this hypothesis, we simulate an unphosphorylated, cytosolic (i.e., no membrane) *ζ*-chain. Surprisingly, we found rate differences also persist without the presence of a membrane. Simulations of a cytosolic *ζ* show reduced preference, in agreement with intuition. However, phosphorylating the sequence 123456 (i.e., N-to-C terminus, membrane-proximal-to-distal, if there were a membrane) is still 2-fold more likely than phosphorylating 654321 (i.e., C-to-N terminus, membrane-distal-to-proximal, if there were a membrane), as shown in Fig. 2 A.ii,B.ii. When we re-simulate cytosolic *ζ*-chain, but now with its six tyrosines spaced evenly along the amino acid sequence, there is no longer any preference for for phosphorylation sequence (Fig. 2A.iii,B.iii). Thus, the emergent preference arises solely from the locations of the tyrosines in *ζ*-chain. The membrane-distal tyrosine is twelve amino acids away from the C-terminus while the membrane proximal tyrosine is twenty-one amino acids away from the membrane. We intuitively understand it as follows. In all cases, the phosphorylation site is attached to a chain of the same length. However, steric hindrance depends on location — specifically, steric hindrance is reduced closer to the chain’s free ends, where there are effectively fewer nearby amino acids to get tangled. These results suggest that changing binding site locations by as little as 10 amino acids is sufficient to induce a preferential binding sequence.

### Local stiffening

One way in which disordered proteins can participate in signaling cascades is to undergo a disorder-to-order transition (8, 22, 45) upon post-translational modification (e.g., tyrosine phosphorylation), becoming locally stiff, as shown schematically in Fig. 1Bii.

Previous experimental-modeling work (19, 20) suggest that the tyrosines on the *ζ*-chain experience phosphorylation rates that were enhanced by previous phosphorylations. Can this be explained by phosphorylation-induced local stiffening?

#### Local stiffening modulates binding rates to multi-site disordered domain

In Fig. 3A, we compute the average binding rate of a kinase to the membrane-bound *ζ* domain in its unphosphorylated state. Here, each average binding rate is weighted by the most likely (de)phosphorylation sequences. (Average binding rates unweighted by their path’s probability are shown in Supporting Material Fig. S3.) Without disordered-to-ordered transitions, the average kinase binding rate decreases as more phosphorylations occur. Because there is a natural preference to phosphorylate membrane distal-to-proximal, it is most likely that the membrane-proximal tyrosine will be the last tyrosine phosphorylated. This tyrosine has a lower binding rate, bringing down the average binding rate of the last phosphorylation compared to the first event, when the faster, membrane-distal sites dominate.

However, if phosphorylation introduces local stiffening, then a single phosphorylation event creates an overall increase in the average binding rate of the kinase to another tyrosine. This effect increases with total phosphorylations and degree of local stiffening per phosphorylation. For example, if each phosphorylation locally stiffens 11 amino acids (∼ 1/12 of the polymer length) then the sixth kinase binding event will occur almost two times times faster than the first (larger range shown in Supporting Material Fig. S4). In agreement with on-lattice simulations (22), local stiffening causes the polymer to be more elongated on average and sample more configurations where the remaining tyrosines are kinase-accessible. Through this mechanism, disordered-to-ordered transitions can increase the binding rate of a kinase to the remaining unphosphorylated tyrosines.

Interestingly, there is a decrease in the average binding rate between the fourth and fifth binding event. We explain this phenomenon, influenced by the second-last tyrosine Y83, in Supporting Material Fig. S5.

While local stiffening at phosphorylated tyrosines enhances the binding rate of kinases to unphosphorylated tyrosines, it could also enhance the binding rate of phosphatases to phosphorylated tyrosines. We therefore explore how dephosphorylation (i.e., loss of local stiffening) impacts the binding rate of phosphatases to the domain in Fig. 3B. For example, if each phosphorylation locally stiffens 11 amino acids, then we assume each dephosphorylation similarly relaxes 11 amino acids. As dephosphorylations occur, entropic flexibility is returned to the polymer. By the sixth dephosphorylation event, the average phosphatase binding rate is decreased by half.

#### Binding rate cooperativity creates ultrasensitivity, even in reversible symmetric phosphorylation cycles

To simulate reversible phosphorylation cycles, we explore two models of dephosphorylation: 1) constant dephosphorylation, in which the phosphatase enzymatic domain is assumed to be small enough that steric effects are insignificant, and 2) steric dephosphorylation, in which the phosphatase enzymatic domain is large enough that the steric effects we report above significantly influence dephosphorylation.

In the first model, the phosphatase is assumed to be of negligible size, and therefore dephosphorylation occurs at a constant rate at each site. Using the site-specific binding rates determined above for membrane-bound *ζ*, we calculate the steady state fraction of sites phosphorylated, with dose-response curves shown in Fig. 4A. As expected, when no local stiffening occurs, reversible phosphorylation has approximately Michaelis-Menten kinetics. However, as the number of amino acids stiffened per phosphorylation increases, the dose-response curves become steeper. This ultrasensitivity can be quantified as a Hill coefficient, here defined as the maximum logarithmic slope of each curve (see Supporting Material text and Fig. S6 for definition of Hill coefficient), attaining a Hill coefficient of 1.8 when approximately 1/12 of the chain is stiffened per phosphorylation event. Therefore, the binding rate cooperativity introduced by local disordered-to-ordered transitions can lead to ultrasensitivity in a signaling network.

**Figure 4:**
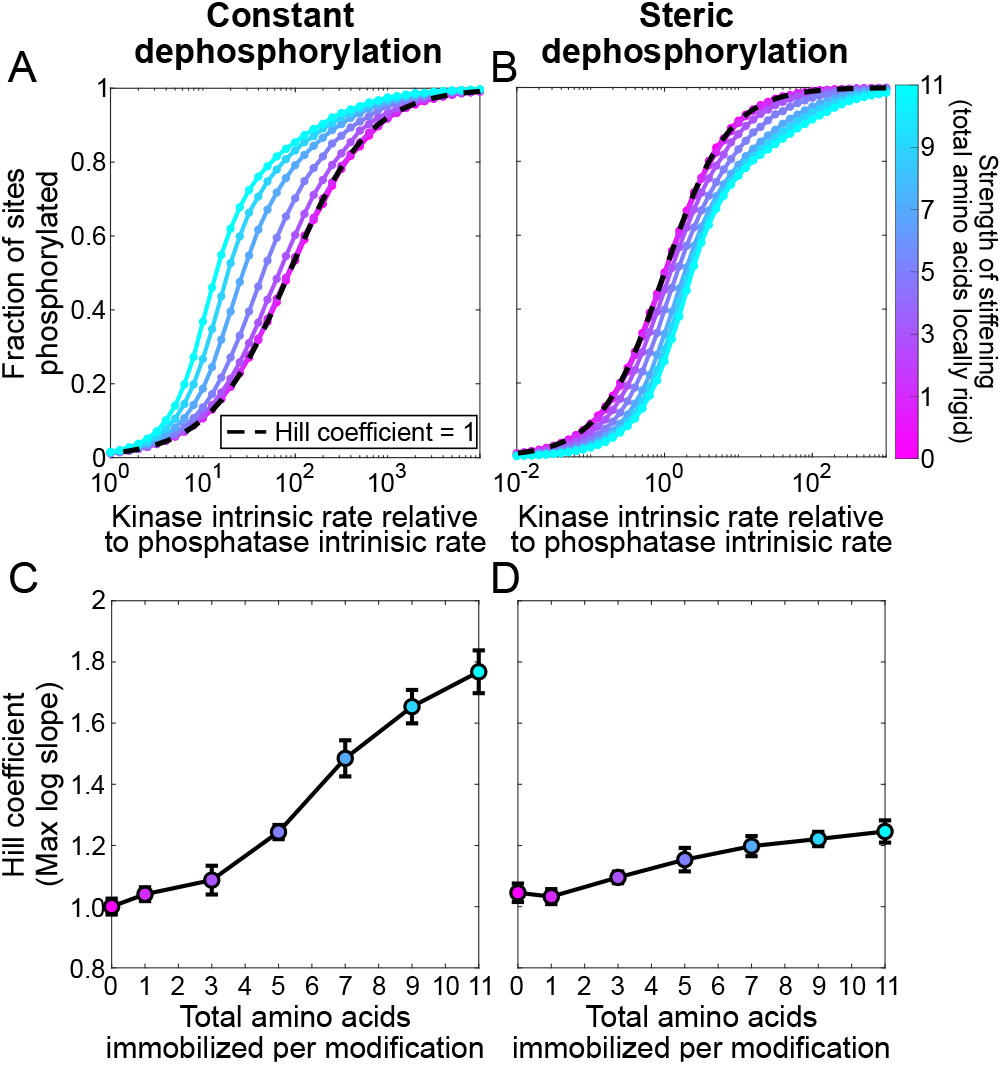
Emergent cooperativity from binding rate enhancement implies switch-like dose-response, even in symmetric phosphorylation/dephosphorylation cycles. (A,B) Fraction of sites phosphorylated as a function of the kinase-to-phosphatase activity ratio (measured by the ratio of free-space rates) for different ranges of local stiffening (colorbar), assuming a phosphatase with (A) negligible size or (B) radius of 2.1 nm, equal to the kinase. Black dashed line indicates a linear dose-response, i.e., with Hill coefficient 1. (C,D) Hill coefficients for different ranges of local stiffening per phosphorylation event, assuming a phosphatase of (C) negligible size or (D) radius of 2.1 nm. Error bars indicate root-mean-square error from a cubic polynomial fit to slope.

**Figure 5:**
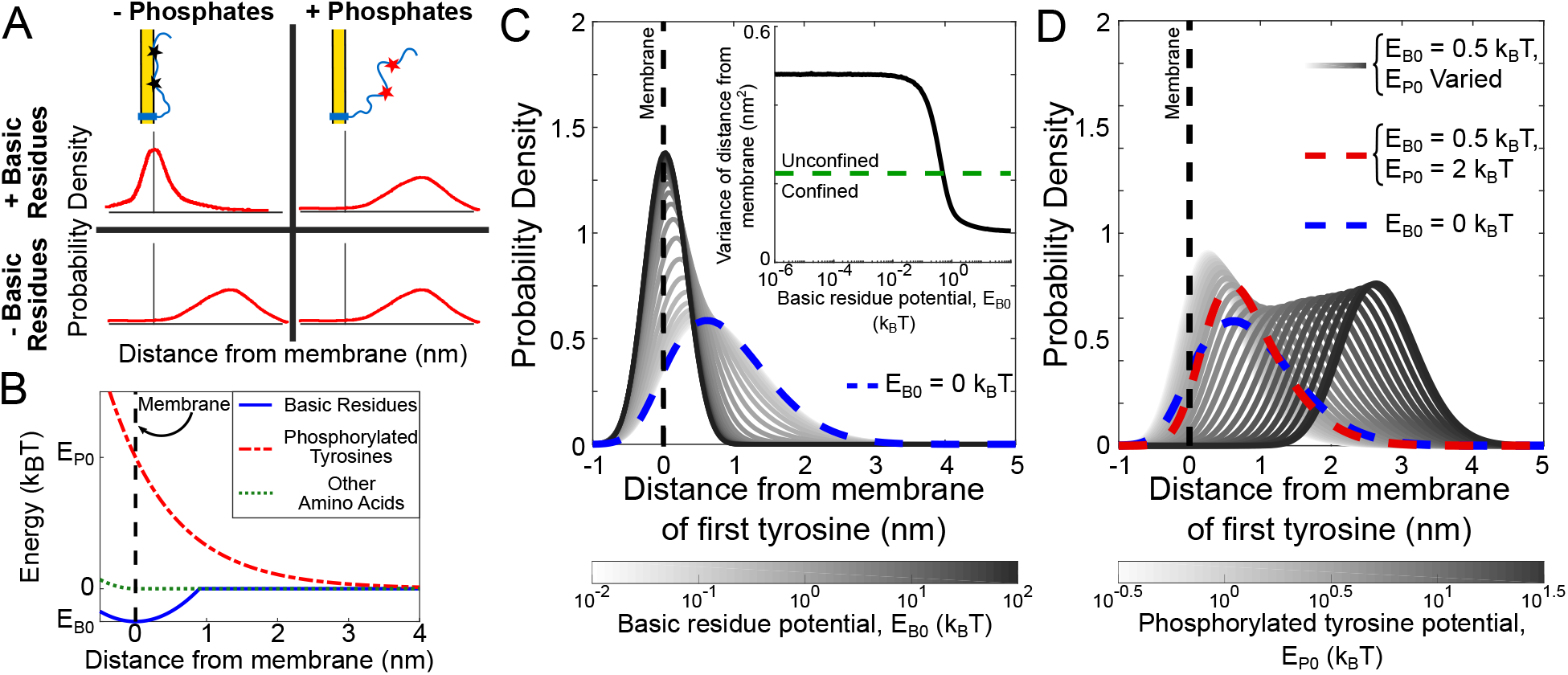
Simplified membrane interaction model sets constraints on the strength of basic residue attraction and phosphorylation-driven repulsion from membrane. (A) Schematic diagrams reflecting probability density of tyrosines under different conditions. Based on previous experimental studies, our model must be consistent with the following: Probability density of tyrosines is narrow and close to the membrane for wild type *ϵ*, but widens and shifts away from the membrane when it is phosphorylated or the basic residues are mutated. (B) Interaction potentials used in simplified model. Basic residues experience an attractive potential of depth *E*_B0_ near the membrane and experience zero potential one Debye length (∼ 1nm) away from the membrane (blue solid line). Phosphorylated tyrosines experience a repulsive potential with strength *E*_P0_ (red dashed line). Unphosphorylated tyrosines and all other amino acids are hindered from entering the membrane but otherwise experience no potential (green dotted line). (C) Probability density of the distance from the membrane of the 1st tyrosine of *ϵ*, assuming no phosphorylated tyrosines, for varying strengths of the basic residue potential, *E*_B0_ (white: low; black: high). The tyrosine moves close to the membrane as *E*_B0_ is increased. Probability density when there is no basic residue potential (*E*_B0_ = 0 *k*_B_*T*) is shown as blue dotted line. (Inset) Variance of probability density of 1st tyrosine over range of basic residue potential strengths. Green dashed line shows the characteristic *E*_B0_ = 0.5*k*_B_*T* required to confine the tyrosine to the membrane, defined in the text. (D) Probability density of the location of 1st tyrosine assuming all tyrosines are phosphorylated, for *E*_B0_ = 0.5 *k*_B_*T* (the value required to confine the tyrosine to the membrane assuming no tyrosines are phosphorylated), and varying phosphorylated tyrosine potential strength, *E*_P0_ (low white, high black). Probability density when *E*_P0_ = 2 *k*_B_*T* is shown as red dashed line, reflecting the *E*_P0_ value needed to approximately return to the distribution when *E*_B0_ = 0 *k*_B_*T* (blue dashed line).

In the second model, the phosphatase is assumed to experience the same steric constraints as the kinase. This model is reasonable since we estimate the size of the phosphatase domain of CD45 to be 3.4 nm. We find dose-response curves shown in Fig. 4B. At 1/12 chain local stiffening per phosphorylation, the Hill coefficient is 1.3. Therefore, even under steric dephosphorylation, a reversible system with local disordered-to-ordered transitions is still capable of creating mild ultrasensitivity. Below, in Fig. 9, we show that if ZAP70 protects phosphorylated tyrosines from dephosphorylation, ultrasensitivity is further enhanced, and is still strong when dephosphorylation experiences the same steric enhancement.

Note by “intrinsic rate”, we mean 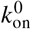, which is the free-space (solution) rate if the reaction did not experience any occlusion. In Fig. 4A, the kinase activity is challenged by significant steric hindrance (but not the phosphatase), so, much more kinase is required to overcome a given phosphatase concentration compared to Fig. 4B.

### Membrane affinity

Recent evidence (24, 46) suggests a model for T Cell signaling in which, before receptor triggering, the E- and *ζ*-chains are unphosphorylated and membrane-associated, protecting them from phosphorylation by kinases (Fig. 1Biii), only dissociating from the membrane after triggering, at which time they can become phosphorylated. But TCR phosphorylation is one of the first steps in triggering (23), raising a “chicken-and-egg” the question: how are the first chains phosphorylated?

A possible hypothesis (27, 47–49) to resolve this puzzle is that, prior to triggering, the chain is biased towards the membrane but spends a small-yet-significant time in the cytoplasm, accessible to kinases. Then, upon initial phosphorylation, the bias is shifted towards the cytoplasm, allowing further phosphorylation. This hypothesis suggests a delicate balance between membrane affinity and phosphorylation. In this section, we ask, if phosphorylation controls membrane association, what are the quantitative constraints on the interaction strengths between the chain, the membrane, and the kinase? And what proportion of states are accessible to the kinase before and after initial phosphorylation?

#### Simplified membrane interaction model can explain both basic residue and phosphorylation effects

Experimental data suggests that *ϵ* membrane-association prior to TCR triggering has two features: first, that the basic residues in *ϵ* are required for membrane association, and second, that fully phosphorylated *ϵ* does not associate with the membrane. We create a simplified electrostatic interaction model, where phosphorylated tyrosines feel a repulsion from the cell membrane and basic residues feel an attraction, shown in Fig. 5B and described by Eq. 5-7. Each potential is parameterized to reproduce the two experimental phenomena. We first look at how strong the basic residue-membrane attraction, *E*_B0_, needs to be in order to confine the tyrosines to the membrane, as defined by the horizontal line in Fig. 5C, inset. We find that the minimum *E*_B0_ for basic residues to achieve this is 0.5 *k*_B_*T* In other words, this is the energetic cost to overcome the chain’s entropic pull away from the membrane.

We next tune the strength of the phosphorylation potential, *E*_P0_, such that tyrosine phosphorylation compensates for the basic residues (i.e. to match the distribution where *E*_B0_ = 0), shown in Fig. 5D. A priori, we expected that in order for full phosphorylation to be enough to counteract the membrane association created by the basic residues, we would require that

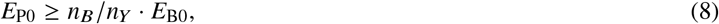

where *n*_*B*_ and *n*_*Y*_ are the numbers of basic residues and tyrosines, respectively. Indeed, in Supporting Material Fig. S7, we find that for the *ζ* chain, this is consistent with simulation. However, for *ϵ*, our simulations indicate that *E*_P0_ ∼ 2*k*_B_*T* is sufficient for its two tyrosines to compensate for the basic residues (Fig. 5D), in contrast to Eq. 8, which predicts a minimum *E*_P0_ of 3.5*k*_B_*T* We can explain this by examining the distribution of basic residues compared to tyrosines. For *ζ*, there are both tyrosines and basic residues along the full length of the domain. In *ϵ*, the two tyrosines are clumped at the membrane-distal end of the domain, with fewer basic residues nearby.

#### Phosphorylation-modulated membrane association allows some early phosphorylation, and at least a small acceleration for late phosphorylation

Using the parameter constraints found above, we examine how phosphorylation influences binding kinetics for *ϵ*, shown in Fig. 6A. At the minimum *E*_P0_ = 2*k*_B_*T* the average kinase binding rate increases approximately 10% between the first and the second phosphorylation. When *E*_P0_ is stronger (*E*_P0_ ≥2*k*_B_*T*), the average kinase binding rate increases significantly for future phosphorylation events (up to 50% for *E*_P0_ = 10*k*_B_*T*).

**Figure 6:**
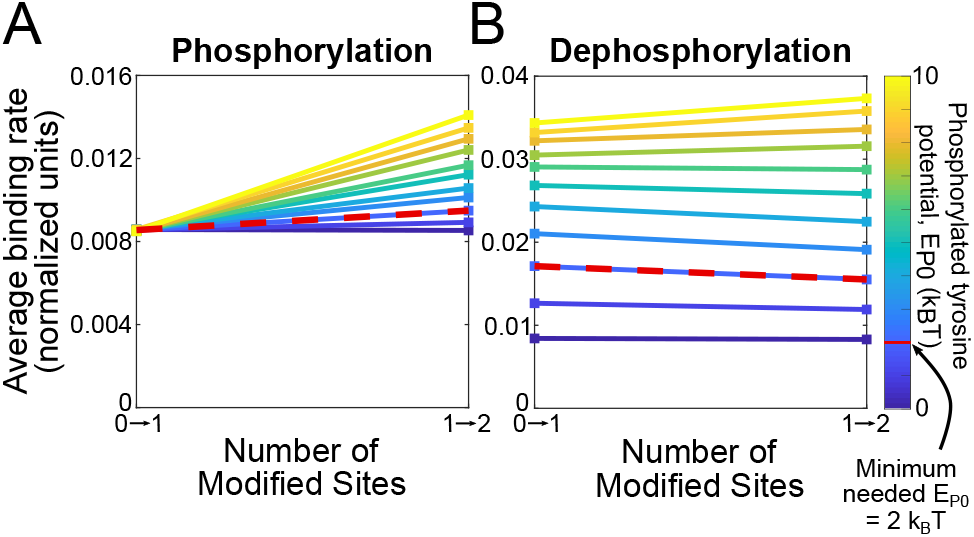
Phosphorylation-driven modulation of membrane association allows tyrosines to operate as a regulatory switch yet still remain accessible in the “off” state. Average binding rates of (A) kinase and (B) phosphatase binding to *ϵ* at different (de)phosphorylation states for varying strengths of phosphorylated tyrosine potential (*E*_P0_) (blue: weak; yellow: strong). Both kinase and phosphatase are assumed to have radius 2.1 nm. Rates are normalized to the free-space binding rate.

We next investigate dephosphorylation in Fig. 6B. Interestingly, there is a weak increase in dephosphorylation rates, in contrast to the stiffening model above. We understand this as follows. As more residues become dephosphorylated, more sections of the chain become restricted to the membrane. The remaining phosphorylated tyrosines remain far from the membrane by the repulsive phosphate-membrane interaction, and therefore more accessible to the phosphatase. The cooperative effect of dephosphorylation only produces a small increase (1.15-fold) in binding rate since the membrane still creates a large steric barrier to phosphatase binding.

Investigation of the reversible system, shown in Supporting Material Fig. 8, shows that stronger phosphate-membrane interactions (*E*_P0_ > 2*k*_B_*T*) cause the dose-response curve to diverge slightly from Michaelis-Menten kinetics under both constant and sterically-influenced dephosphorylation.

### Multiple simultaneous binding

One of the first molecular participants in signaling downstream of TCR is ZAP70, which binds phosphorylated tyrosines (18, 19, 29, 50, 51). A recent in vivo experimental study found that approximately six ZAP70 molecules are bound per TCR (29), despite having ten possible binding sites. This raises two questions: Do entropic constraints arising from polymer flexibility prevent more ZAP70 ligands from binding (in addition to possible low occupancy set by dissociation constant)? And, if so, are there advantages to the cell in not using the full array of possible ZAP70 binding sites? To address this question, we simulate binding of a ligand, ZAP70, to the full TCR, when other ligands are already bound, shown schematically in Fig. 1Biv.

#### Multiple binding to multi-site disordered domain gives rise to negative cooperativity

We calculate the average binding rates of ligands to all 6 membrane-bound chains of TCR, assuming a given number of previously bound ligands. In this section, rather than a single chain, we simulate all 6 chains anchored to the membrane (38), and rather than represent all tyrosines as individual sites, we place a binding site at the center of each ITAM. In other words, in the previous sections, we independently treated Lck and CD45 enzymatic domains interacting with single tyrosines, whereas here we treat ZAP70 interacting with two tyrosines on an ITAM.

First, we ran simulations to see if there is a hard limit to the number of ZAP70 molecules that can bind to the TCR. We find that at our estimate for the size of ZAP70, there are configurations that allow binding a full ten ZAP70 molecules to the 6 chains. Indeed, ten molecules with radius of up to 6.9nm (larger than our estimate of the size of ZAP70) can still fit, as shown in Fig. 7B. Given that the steric limit is not being reached, is there a limit set by entropic effects?

**Figure 7:**
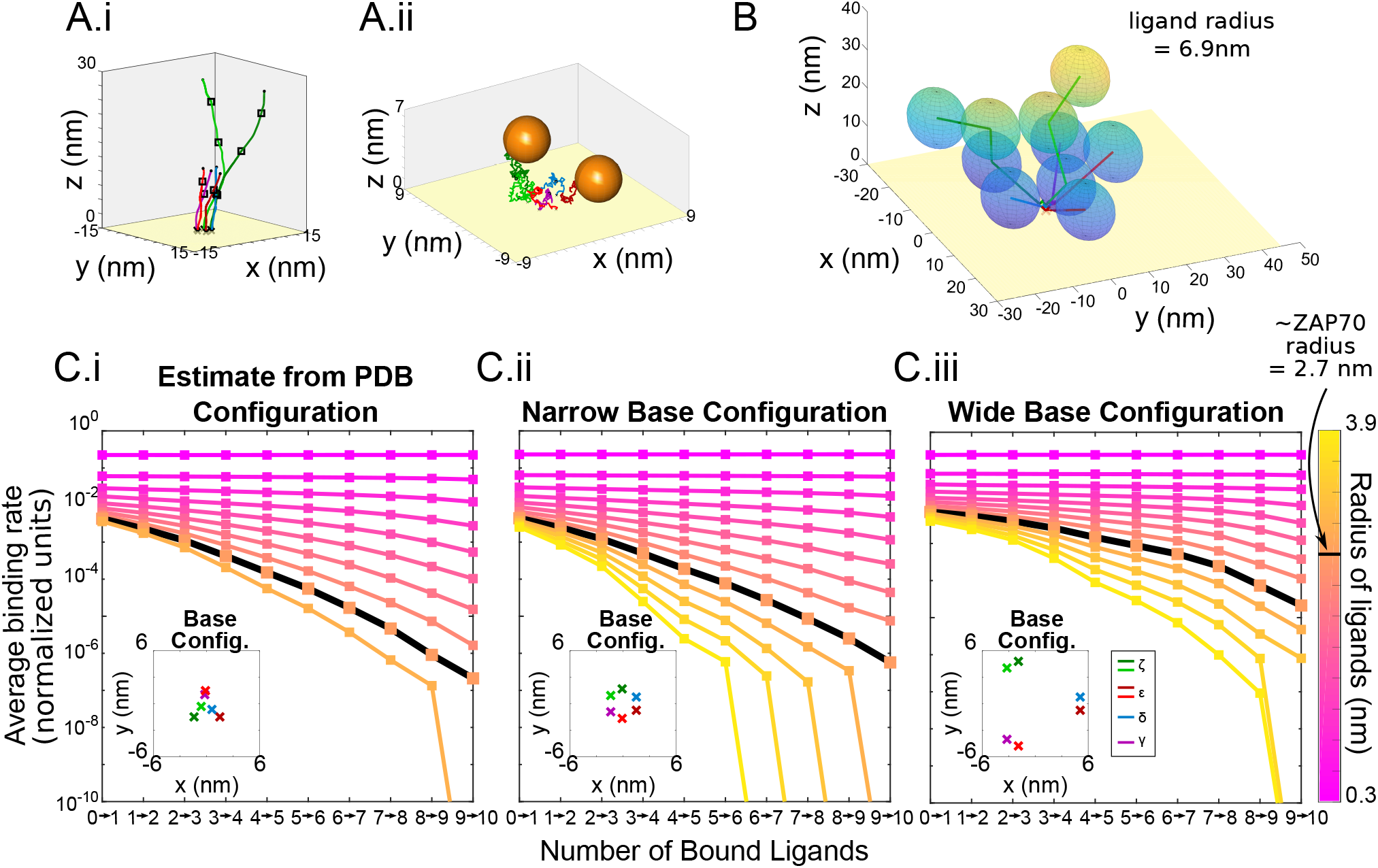
Simultaneously-bound disordered regions have a reduced binding rate as they become more crowded, even before the steric limit. (A) Snapshots of simulated TCR subunits and binding sites (black squares) (Ai) in extended conformation and (Aii) at equilibrium with two bound ligands. (B) Simultaneous binding of full TCR is not sterically prohibited at physiological ligand radii. View of single TCR configuration with ten bound ligands where each ligand has radius of 6.9 nm. (C) Average binding rates to TCR against number of ligands bound for varying ligand size (color bar; pink: small; blue: large) and TCR subunit configuration (Ci) Estimated from PDB structure of TCR, (Cii) 1.5nm, and (Ciii) 5nm.

**Figure 8:**
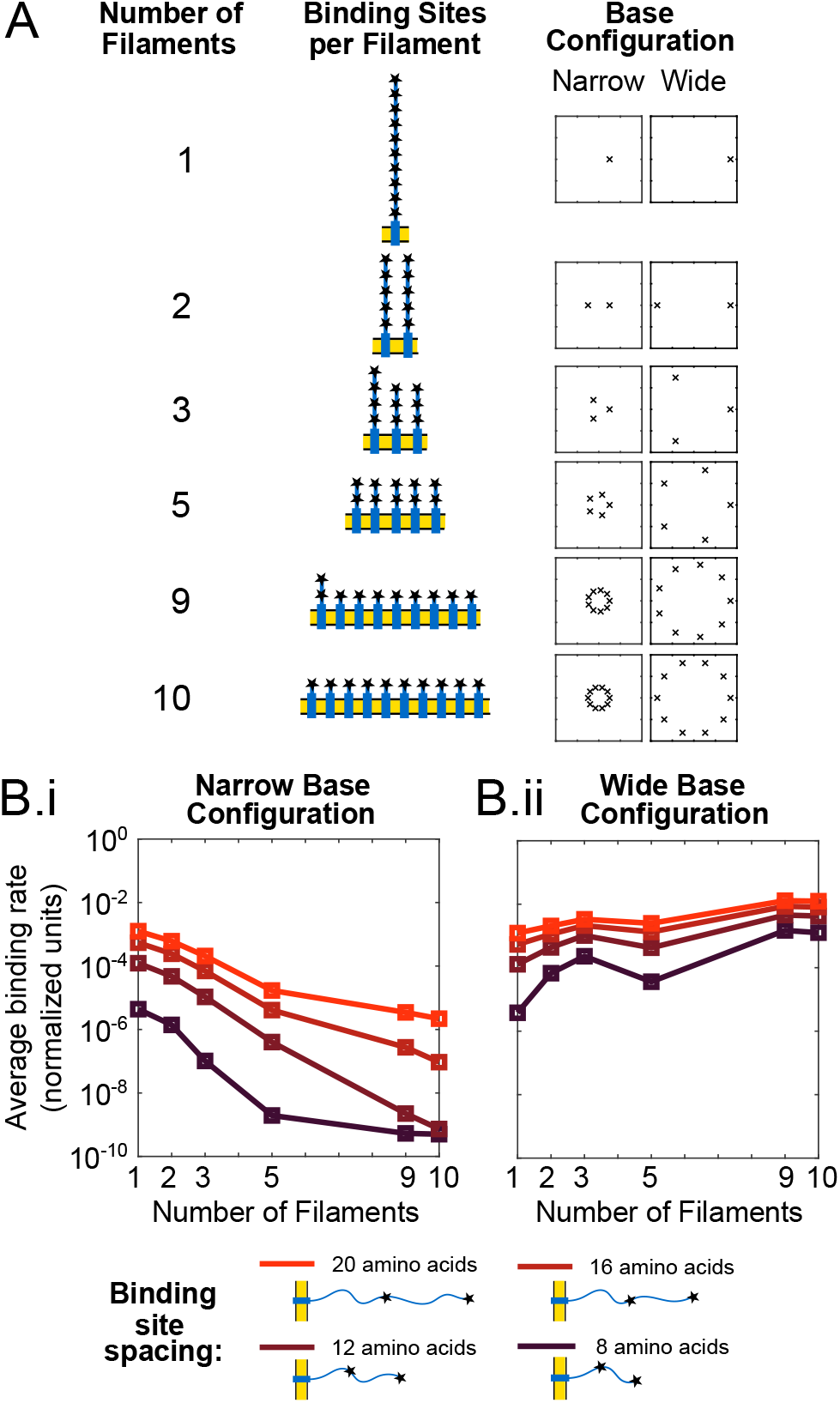
Optimal distribution of ten binding sites on multiple chains is dependent on membrane spacing of subunits. (A) Schematic for distribution of 10 binding sites on multiple chains. For each, chains are distributed evenly on a (1) narrow circle of radius 1.5 nm, and (2) wide circle of radius 5 nm. (B) Average path-weighted binding rates of sixth binding event to constructed domain against number of chains in domain (color bar; dark red: short spacing between binding sites; bright red: long spacing) and subunit configuration (Bi) 1.5nm radius, (Bii) 5nm radius. Simulations explore different spacings between binding sites from 8 to 20 amino acids, indicated by color. Ligand radius is 2.7 nm.

**Figure 9:**
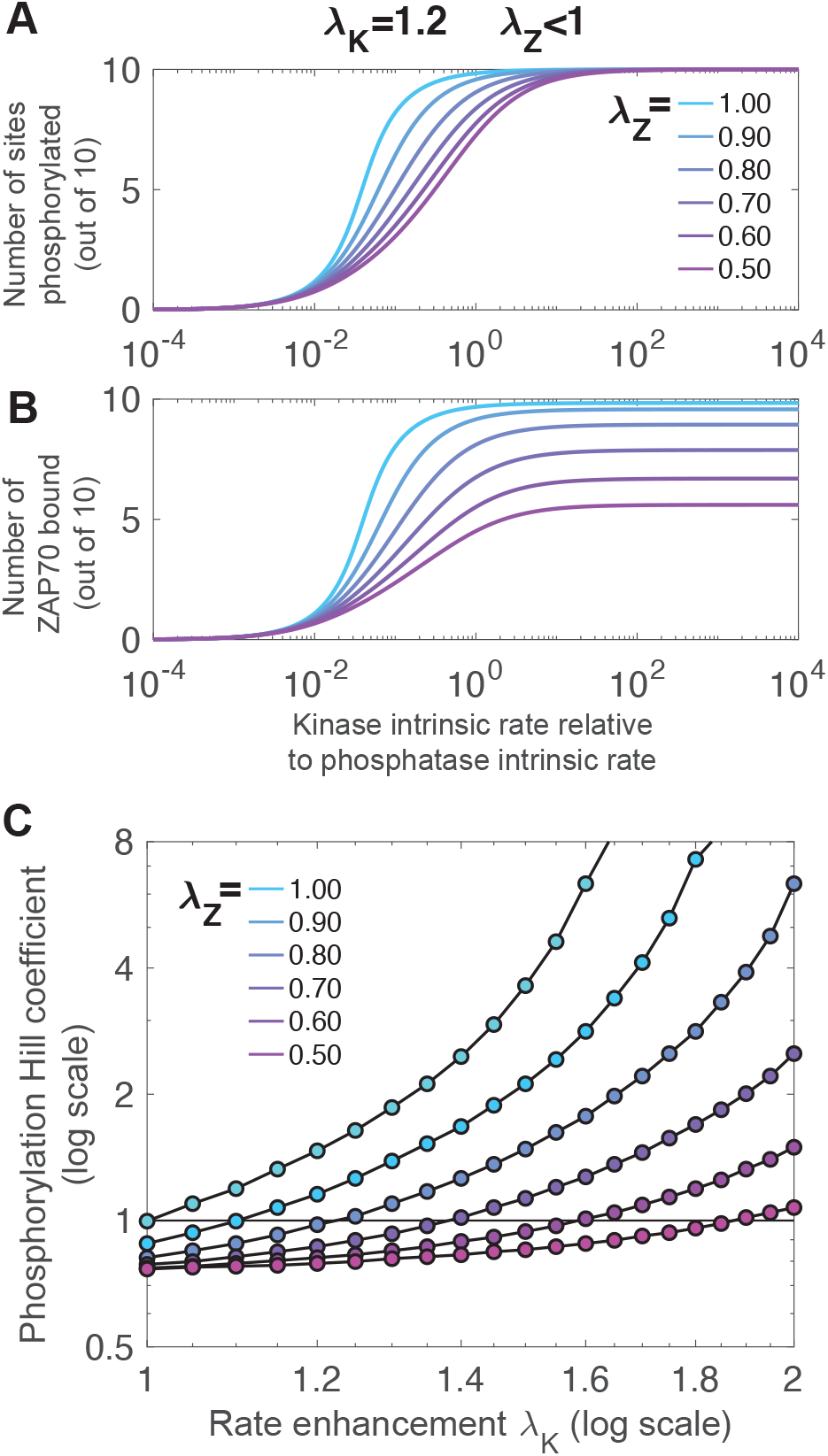
Integrative model shows counteracting effects of rate enhancement and rate reduction. Rate reduction from ZAP70 binding (λ_*Z*_ < 1 indicated by color) abrogates the switch-like dose response conferred by rate enhancement of kinase phosphorylating TCR λ_*K*_ = 1.2 for both amount of phosphorylation (A) and amount of ZAP70 bound (B), in addition to reducing the saturating value (*E*_max_) of ZAP70 binding (B). (C) Ultrasensitivity quantified using the log-effective-concentration definition of Hill coefficient (Supporting Eq. 2) for a range of λ_*K*_ ≥ 1 and λ_*Z*_ ≤ 1.

As more ligands are simultaneously bound to the TCR, the average binding rate decreases. For a ligand with size of ZAP70, approximately 2.7 nm radius, the binding rate for the seventh ligand is approximately 200 times lower than the binding rate of the first ligand (Fig. 7Ci). Simulations where the chains are anchored further apart, as shown in Fig. 7Ciii, exhibit significantly less hindrance.

In both cases, the rate decrease implies a “negative cooperativity” effect, which could allow TCR to recruit a regulated number of ZAP70 molecules while still maintaining high binding rate. The first few ZAP70 molecules are able to bind to the domain relatively easily, but it quickly becomes prohibitive to bind more. The alternative way of regulating a limit around 6 ZAP70 molecules would be to have only 6 binding sites, but then the total binding rate would be lower (since there would be 4 fewer opportunities to bind).

#### Optimal distribution of binding sites on multiple nearby chains

The above finding suggests that the chain’s entropy influences how readily a multi-chain receptor can become loaded with signaling molecules. This led us to ask: Given ten binding sites, if the goal is fast loading, is it preferable to have 10 different chains with 1 binding site each, or 1 chain with all 10 binding sites (or, perhaps, have the 10 sites distributed on 6 chains, as they are for TCR)? A priori, it could be that the two effects of a nearby membrane and ligands bound to neighboring chains “in trans”, together, prohibit binding, leading to a preference for concentrating the sites on fewer chains. Alternatively, it could be that other molecules bound “in cis” on the same chain prohibit binding, leading to a preference for sites that are distributed across many chains.

We simulate 10 sites (for a ligand approximately the size of ZAP70) distributed on 1, 2, 3, 5, 9 or 10 chains, as shown in Fig. 8A. The chains are assumed to each be anchored to the membrane on the perimeter of a circle. The radius of the circle is either 1.5nm, similar to the spacing found in the crystal structure of the TCR (38), or wider at 5nm. We report the average rate of the 6th binding event.

We find that, if the chains are in the narrow configuration, fastest binding occurs when the sites are all on a single chain (Fig. 8Bi). This result holds over a wide range of spacings between the sites on the chain. When the chains are anchored further apart, we find that the fastest binding is achieved when the ten sites are distributed across ten chains (Fig. 8Bii), in agreement with intuition. These results suggest a consideration for engineered receptors: There are significant rate differences obtained by having the same signaling modules on multiple versus single chains. Specifically, for the base separation distance estimated for TCR, we find that it is beneficial to have the sites on the same chain to minimize neighbor interference.

For these simulations, we explore different spacings between the tyrosines. For simplicity we assume these sites are evenly spaced, unlike the real TCR, with spacing ranging from 8 to 20 amino acids, indicated by color in Fig. 8. The relative performance of different filament distributions do not depend on within-chain spacing, although overall steric hindrance increases with less spacing, as intuitively expected.

For a receptor with 10 binding sites, the sixth binding event can happen in 1260 ways. Interestingly, we find that the rates of these 1260 configurations can have multimodal distributions. In Supporting Material Fig. S11, we explore how this multimodality arises.

### Integrative model of nonlinear modules

The models above each lead to nonlinear behavior in different modules of the TCR triggering process. To roughly summarize: First, for the tyrosines on a single *ζ* chain or *ϵ* chain, phosphorylation-induced stiffening or membrane affinity control lead to positive rate enhancements for the accessibility to kinases and/or phosphatases. And second, for the full 6-chain TCR, multiple simultaneous binding by ZAP70 leads to a rate decrease for subsequent accessibility. So the question arises, what is the net effect?

We explore this question in an integrative model combining phosphorylation, dephosphorylation and ZAP70 binding of all 10 sites on a TCR. We coarsen the modeling approach and make the simplifying assumption that rate modifications act similarly independent of the site. This allows a continuous-state model expressed as a system of nonlinear differential equations. We assume that of *N* = 10 sites, *n*_*P*_ are phosphorylated, and *n*_*Z*_ are ZAP70-bound. Only phosphorylated sites can be ZAP70-bound, and ZAP70 sites are protected from dephosphorylation (which is a simplification of the tandem ZAP70-ITAM interaction (17)). Therefore,

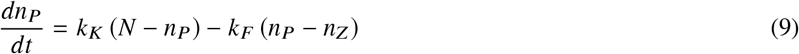

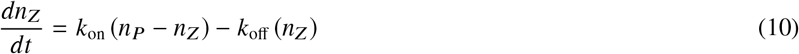

where *k*_*K*_ *k*_*F*_ *k*_on_, and *k*_off_ are the rate of phosphorylation, dephosphorylation, ZAP70 binding, and ZAP70 unbinding respectively. The nonlinear modules that emerged in previous sections of this paper represent non-constant rates, which we approximate from the previous sections by fitting to exponential forms

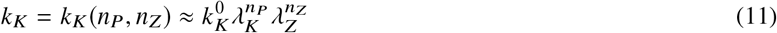

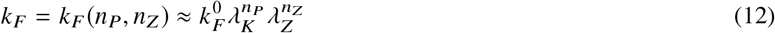

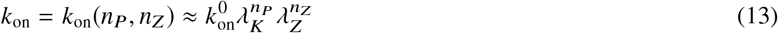

where λ_*K*_ > 1 is the rate enhancement, either from stiffening or membrane dissociation, and λ_*Z*_ < 1 is the rate decrease from multiple ZAP70 binding. For kinetic rates, *k*_off_ ≈0.2 s^−1^ (17), and we sweep over a range of *k*_*K*_ and *k*_*F*_ to produce dose-response curves. For the nonlinearities, we estimate λ_*K*_ ≈ 1.2 for stiffening from Fig. S3, and similarly λ_*K*_ ≈ 1.2 from Fig. S7. In other words, each phosphorylation increases the subsequent phosphorylation by about 20%. We estimate λ_*Z*_ ≈ 0.6 from the rate reductions shown in Fig. 7. Code is available at github.com/allardjun/EntropicMultisiteIntegrative (DOI:10.5281/zenodo.4118734).

#### Counteracting effects of rate enhancement and rate reduction

Dose-response curves for various combinations of λ_*K*_ and λ_*Z*_ are shown in Fig. 9 and Fig. S12.

First, we simulate in the absence of steric inhibition from multiple ZAP70 binding (λ_*Z*_ = 1). We find that when ZAP70 is present and protects phosphorylated sites from dephosphorylation, then even with symmetric phosphorylation and dephosphorylation, the rate enhancement λ_*K*_ > 1 leads to significant ultrasensitivity. This is related to a known phenomenon where sequestration of sites gives ultrasensitivity (52). In addition to ultrasensitivity, the rate enhancement effect reduced the EC50, in other words, increases potency.

Next, we include the steric inhibition from multiple ZAP70 binding (λ_*Z*_ < 1). The multiple binding rate reduction reduces the saturating number (*E*_max_) of ZAP70 binding, but not phosphorylation. Depending on *k*_off_, this leads to steady state of ∼ 6 ZAP70 per TCR, even though the rate reduction is not a sharp cutoff. In agreement with intuition, this occurs around when 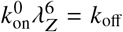.

The rate reduction of ZAP70, when λ_*K*_ = 1 (no rate enhancement), leads to shallow dose-response curves. For this reason, in Fig. 9C, we quantify the ultrasensitivity using the log-concentration definition (see Supporting Eq. S2) which can be negative to indicate shallow responses. When both λ_*K*_ > 1 and λ_*Z*_ < 1, the effects counteract each other. Roughly, an enhancement of λ_*K*_ = 2 is required to counteract a reduction of λ_*Z*_ = 0.5.

At λ_*Z*_ = 0.6 and λ_*P*_ = 1.2, the reduction effect dominates, leading to a negative Hill coefficient. Interestingly, in previous work (19), we found a lack of ultrasensitivity in the presence of multiple ITAMs. Note that at these parameters, the rate enhancement still leads to increased potency — this is not abrogated by the multiple-binding-induced reduction.

## DISCUSSION

Multisite modification of signaling molecules leads to high combinatorial complexity that is challenging to study. Since there are 10 ITAMs on a TCR, we were required to simulate ∼1000 (= 2^10^) binding states and ∼3.6 million (= 10 factorial) possible sequences of phosphorylation. Other explorations in this work have similar combinatorial complexity, for example, the 6 tyrosines on CD3*ζ* imply 64 binding states and 720 sequences. Furthermore, a major opportunity of computational simulation is its ability to simulate counter-factual parameters, for example larger or smaller ligand radii, and ITAMs arranged on different numbers of chains, since these counter-factuals provide insight into the real parameters. The coarse-grain modeling approach we use here — representing structured domains as simple rigid bodies and disordered regions as simple polymers, and membrane interactions with simple potentials — are computationally efficient enough for us to perform such parameter exploration.

Previous modeling work on TCR largely falls in two categories. First are models of the kinetics of signaling by TCR, in which the system is modeled as transitions between distinct molecular states (e.g., membrane-bound or not (27)), for example using differential equations or Markov chains to model binding to multisite molecules (53–57). Second, several detailed molecular models have focused on the TCR’s transmembrane and extracellular domains (e.g., see (58, 59)), due to the challenges of simulating the high-intrinsic disorder of the chains (60, 61) – with notable exception of (43). Our work attempts to bridge this gap, connecting sub-molecular states with the kinetics that provide inputs for models of the signaling cascade. While the coarse granularity omits many details, such as sequence-dependent persistence lengths (62), we join a growing body of modeling efforts (63, 64) at the granularity between particles representing individual proteins and particles representing individual atoms, which has been particularly fruitful for intrinsically disordered domains (10–12, 22, 65)

Multi-site reactions can be classified in two categories: sequential or random. For CD3*ζ*, evidence for sequential phosphorylation is mixed (54, 66, 67). Our work offers a possible resolution. As opposed to obligate sequential modification, meaning the previous event is required before the next event can occur, we find that multi-site disordered domains can display preferential sequential modification, meaning that previous events reduce the probability of future sequences, but do not prohibit them. Note that the extensive theoretical studies of the consequences of sequential binding (21, 53–56) assumed obligate sequences. In some cases, the sequence is assumed, and the mechanism enforcing it is unclear. In contrast, in our current model, the sequence is an emergent property of the distribution of sites along the chain. We find that the membrane increases the differences in binding rate and therefore amplifies the preference, but it is the positions of the tyrosines along the chain that are the primary drivers.

Previous results suggested that the *ζ*-chains exhibit cooperative phosphorylation (19). We demonstrate that phosphorylation-induced local structuring of *ζ* naturally leads to cooperativity, offering a possible explanation for these results. Alternative models to explain the TCR ultrasensitivity include TCR clustering (68) and changes in lipid composition (49, 69, 70). Thus, our model suggests the following experiments: If it is local structuring that drives cooperativity, adding extra residues, increasing the spacing between ITAMs would reduce ultransensitivity, while reducing the number of residues between ITAMs would increase ultrasensitivity.

Prior to T cell triggering, the CD3*ϵ* domains of the TCR associate with the cell membrane, with the tyrosines primarily embedded in the bilayer (24–26). Post-triggering, they dissociate from the membrane, revealing the tyrosines for phosphorylation (26). Thus, membrane association could act as a switch controlling the activity state of the CD3*ϵ* chain. However, given that tyrosine phosphorylation is one of the first events in TCR triggering, this hypothesis features a “chicken-and-egg” paradox. A plausible resolution (26) is that the unphosphorylated chain is exploring an ensemble of configurations in which its accessibility to kinases is limited but not totally forbidden. The question, then, is whether this accessibility changes enough between unphosphorylated and semi-phosphorylated states to provide a switch. We find that, to be consistent with the two observations that both basic residues and phosphostate control membrane proximity, phosphotyrosines in CD3*ϵ* must experience a repulsive potential of approximately *E*_*P*0_ ≈ 2*k*_*B*_*T* However, at this repulsion energy, the semi-phosphorylated state is only ∼10% more accessible than the unphosphorylated state. To get an increase in accessibility of 50% would require *E*_*P*0_ ≈ 10*k*_*B*_*T*, which is several times larger than typical residue-membrane energies (44). Our results do not preclude the membrane association switch hypothesis, but do set a requirement for large interaction energies in order for it to hold, for example due to dynamic changes in lipid composition.

As shown in Fig. 7, we find that ten ZAP70 molecules readily fit on a TCR, but that the seventh ZAP70 molecule binds at a rate ∼ 200-fold slower than the first. To clarify the difference between these two forms of steric occlusion, in the first case, we ask whether all microstates (polymer configurations) in the fully-bound macrostate are forbidden, while in the second case, we ask what proportion of microstates are forbidden. Our finding is that a significant proportion of microstates are forbidden, even at ligand sizes where the fully bound macrostate is not forbidden. In other words, entropy limits binding much before volume exclusion forbids it. When we include these entropic effects in a model of both attachment and detachment (Fig. 9), we find a steady-state with 6-ZAP70-per-TCR stoichiometry, as previously reported (29). This occurs even though the entropic effects do not lead to a sharp cutoff in attachment rate.

These steric effects imply negative cooperativity. Negative cooperativity can endow signaling systems with two features: high turnover of ligands even in high ligand concentration (which might be advantageous if ligands are involved in other reactions), and constant signaling activity in low ligand concentration. Similarly, this might reduce the impact of inhibitors, requiring much higher concentrations of inhibitor to completely turn off signaling (71). This leads to a model prediction: If the non-TCR-binding domains of ZAP70 are removed, the model predicts an increase in ultrasensitivity and a higher bound fraction.

Our model predicts that multiple ITAMs on the same chain will bind to ZAP70 faster than the same number of ITAMs on multiple clustered chains, provided the chains have membrane domains with similar relative spacing as the TCR. For larger relative spacing of chains, the relationship flips, and it is better to have ITAMs on multiple chains (Fig. 8B). This is a testable prediction. Furthermore, even without experiments that modify the spacing, our model predicts the fold-change reduction in binding rates for subsequent ZAP70 molecules. The rate of a single ZAP70 binding to an ITAM has been measured (17). We predict that, for example, the 2nd ZAP70 will bind at rate 20% slower (Fig. 7). If realized, this would add to a growing body of work demonstrating the importance of the disordered regions of amino acids *between* catalytically active or post-translationally modified parts of proteins (10, 13).

While we propose several experimental tests of the model (e.g., addition of residues between ITAMs, removal of the non-binding domain of ZAP70), these are mostly positive tests. There is also a negative test: If the freely-jointed chain model is applicable to TCR chains, then replacing residues between ITAMs in any of the chains should have minimal effect on phosphostate, ZAP70 binding, and ultimately on TCR signaling, provided that the number of residues, intrinsic disorder, and electrostatic properties of the residues are preserved.

## AUTHOR CONTRIBUTIONS

LC wrote simulation code, carried out simulations, analyzed the data and prepared the manuscript. OD helped design the research and prepare manuscript. JA designed the research, supervised the project and helped prepare the manuscript.

## ACKNOWLEDGMENTS

This work was supported by NSF CAREER grant DMS 1454739 to JA, NSF grant DMS 1715455 to JA, and NSF grant DMS 1763272 and a grant from the Simons Foundation (594598, QN), and Wellcome Trust grant SRF 207537/Z/17/Z to OD. We thank Sean Lawley (University of Utah) for valuable discussion.

## SUPPLEMENTARY MATERIAL

An online supplement to this article can be found by visiting BJ Online at http://www.biophysj.org.

## REFERENCES

1. Van Der Lee, R., M. Buljan, B. Lang, R. J. Weatheritt, G. W. Daughdrill, A. K. Dunker, M. Fuxreiter, J. Gough, J. Gsponer, D. T. Jones, P. M. Kim, R. W. Kriwacki, C. J. Oldfield, R. V. Pappu, P. Tompa, V. N. Uversky, P. E. Wright, and M. Madan Babu, 2014. Classification of Intrinsically Disordered Regions and Proteins. Chem Rev 114:6589–6631.

2. Tompa, P., and K.-H. Han, 2012. Intrinsically disordered proteins. Physics Today 65:64–65.

3. Gonfloni, S., J. C. Williams, K. Hattula, A. Weijland, R. K. Wierenga, and G. Superti-Furga, 1997. The role of the linker between the SH2 domain and catalytic domain in the regulation and function of Src. EMBO J 16:7261–7271.

4. Kovar, D. R., and T. D. Pollard, 2004. Insertional assembly of actin filament barbed ends in association with formins produces piconewton forces. Proc Natl Acad Sci 101:14725–14730.

5. Romero, S., C. Le Clainche, D. Didry, C. Egile, D. Pantaloni, and M. F. Carlier, 2004. Formin is a processive motor that requires profilin to accelerate actin assembly and associated ATP hydrolysis. Cell 119:419–429.

6. Duchardt, E., A. B. Sigalov, D. Aivazian, L. J. Stern, and H. Schwalbe, 2007. Structure induction of the T-cell receptor ζ-chain upon lipid binding investigated by NMR spectroscopy. ChemBioChem 8:820–827.

7. Keir, M. E., M. J. Butte, G. J. Freeman, and A. H. Sharpe, 2008. PD-1 and Its Ligands in Tolerance and Immunity. Annu Rev Immunol 26:677–704.

8. Bah, A., R. M. Vernon, Z. Siddiqui, M. Krzeminski, R. Muhandiram, C. Zhao, N. Sonenberg, L. E. Kay, and J. D. Forman-Kay, 2015. Folding of an intrinsically disordered protein by phosphorylation as a regulatory switch. Nature 519:106–9.

9. Bah, A., and J. D. Forman-Kay, 2016. Modulation of Intrinsically Disordered Protein Function by Post-translational Modifications. J Biol Chem 291:6696–6705.

10. Goyette, J., C. S. Salas, N. Coker-Gordon, M. Bridge, S. A. Isaacson, J. Allard, and O. Dushek, 2017. Biophysical assay for tethered signaling reactions reveals tether-controlled activity for the phosphatase SHP-1. Science Advances 3:e1601692.

11. Van Valen, D., M. Haataja, and R. Phillips, 2009. Biochemistry on a leash: The roles of tether length and geometry in signal integration proteins. Biophys J 96:1275–1292.

12. Bryant, D., L. Clemens, and J. Allard, 2017. Computational simulation of formin-mediated actin polymerization predicts homologue-dependent mechanosensitivity. Cytoskeleton 74:29–39.

13. Zhang, Y., L. Clemens, J. Goyette, J. Allard, O. Dushek, and S. A. Isaacson, 2019. The Influence of Molecular Reach and Diffusivity on the Efficacy of Membrane-Confined Reactions. Biophys J 117:1189–1201.

14. Smith-Garvin, J. E., G. A. Koretzky, and M. S. Jordan, 2009. T Cell Activation. Annu Rev Immunol 27:591–619.

15. Lever, M., H.-s. Lim, P. Kruger, J. Nguyen, N. Trendel, E. Abu-shah, P. K. Maini, P. A. V. D. Merwe, M. Lever, H.-s. Lim, P. Kruger, J. Nguyen, N. Trendel, and E. Abu-shah, 2016. Architecture of a minimal signaling pathway explains the T-cell response to a 1 million-fold variation in antigen affinity and dose. Proc Natl Acad Sci 113:E6630–E6638.

16. Love, P. E., and S. M. Hayes, 2010. ITAM-mediated signaling by the T-cell antigen receptor.

17. Goyette, J., D. Depoil, Z. Yang, S. A. Isaacson, J. Allard, P. A. van der Merwe, K. Gaus, M. L. Dustin, and O. Dushek, 2020. Regulated unbinding of ZAP70 at the T cell receptor by kinetic avidity. bioRxiv 2020.02.12.945170; doi: https://doi.org/10.1101/2020.02.12.945170

18. Wang, H., T. A. Kadlecek, B. B. Au-Yeung, H. E. Goodfellow, L. Y. Hsu, T. S. Freedman, and A. Weiss, 2010. ZAP-70: an essential kinase in T-cell signaling. Cold Spring Harbor Perspectives in Biology 2:1–17.

19. Mukhopadhyay, H., B. De Wet, L. Clemens, P. K. P. Maini, J. Allard, P. A. Van Der Merwe, and O. Dushek, 2016. Multisite phosphorylation modulates the T Cell Receptor ζ-Chain potency but not the switchlike response. Biophys J 110:1896–1906.

20. Hui, E., and R. D. Vale, 2014. In vitro membrane reconstitution of the T-cell receptor proximal signaling network. Nat Struct Mol Biol 21:133–142.

21. Gunawardena, J., 2005. Multisite protein phosphorylation makes a good threshold but can be a poor switch. Proc Natl Acad Sci 102:14617–14622.

22. Lenz, P., and P. S. Swain, 2006. An Entropic Mechanism to Generate Highly Cooperative and Specific Binding from Protein Phosphorylations. Curr Biol 16:2150–2155.

23. Dushek, O., 2012. Elementary steps int cell receptor triggering.

24. Xu, C., E. Gagnon, M. E. Call, J. R. Schnell, D. Charles, C. V. Carman, J. J. Chou, and K. W. Wucherpfennig, 2008. Regulation of T cell Receptor Activation by Dynamic Membrane Binding of the CD3E Cytoplasmic Tyrosine-Based Motif. Cell 135:702–713.

25. Guo, X., C. Yan, H. Li, W. Huang, X. Shi, M. Huang, Y. Wang, W. Pan, M. Cai, L. Li, W. Wu, Y. Bai, C. Zhang, Z. Liu, X. Wang, X. F. Zhang, C. Tang, H. Wang, W. Liu, B. Ouyang, C. C. Wong, Y. Cao, and C. Xu, 2017. Lipid-dependent conformational dynamics underlie the functional versatility of T-cell receptor. Cell Research 27:505–525.

26. Zhang, H., S.-P. Cordoba, O. Dushek, and P. Anton van der Merwe, 2011. Basic residues in the T-cell receptor ζ cytoplasmic domain mediate membrane association and modulate signaling. Proc Natl Acad Sci 108:19323–8.

27. Yang, W., W. Pan, S. Chen, N. Trendel, S. Jiang, F. Xiao, M. Xue, W. Wu, Z. Peng, X. Li, H. Ji, X. Liu, H. Jiang, H. Wang, H. Shen, O. Dushek, H. Li, and C. Xu, 2017. Dynamic regulation of CD28 conformation and signaling by charged lipids and ions. Nat Struct Mol Biol 24:1081–1092.

28. Chen, X., W. Pan, Y. Sui, H. Li, X. Shi, X. Guo, H. Qi, C. Xu, and W. Liu, 2015. Acidic phospholipids govern the enhanced activation of IgG-B cell receptor. Nat Comm 6.

29. O’Donoghue, G. P., R. M. Pielak, A. A. Smoligovets, J. J. Lin, and J. T. Groves, 2013. Direct single molecule measurement of TCR triggering by agonist pMHC in living primary T cells. eLife 2:e00778.

30. Zhou, H. X., 2001. Loops in proteins can be modeled as worm-like chains. J Phys Chem B 105:6763–6766.

31. Dignon, G. L., W. Zheng, Y. C. Kim, R. B. Best, and J. Mittal, 2018. Sequence determinants of protein phase behavior from a coarse-grained model. PLoS Comp Biol 14.

32. Reeves, D., K. Cheveralls, and J. Kondev, 2011. Regulation of biochemical reaction rates by flexible tethers. Phys Rev E 84:1–12.

33. Kutys, M. L., J. Fricks, and W. O. Hancock, 2010. Monte Carlo analysis of neck linker extension in kinesin molecular motors. PLoS Comp Biol 6.

34. Fischer, H., I. Polikarpov, and A. F. Craievich, 2004. Average protein density is a molecular-weight-dependent function. Protein Science 13:2825–2828.

35. Yamaguchi, H., and W. A. Hendrickson, 1996. Structural basis for activation of human lymphocyte kinase Lck upon tyrosine phosphorylation. Nature 384:484.

36. Hatada, M. H., X. Lu, E. R. Laird, J. Green, J. P. Morgenstern, M. Lou, C. S. Marr, T. B. Phillips, M. K. Ram, K. Theriault, M. J. Zoller, and J. L. Karas, 1995. Molecular basis for interaction of the protein tyrosine kinase ZAP-70 with the T-cell receptor. Nature 377:27–31.

37. Schrödinger, LLC, 2015. The {PyMOL} Molecular Graphics System, Versioñ1.8.

38. Dong, D., L. Zheng, J. Lin, B. Zhang, Y. Zhu, N. Li, S. Xie, Y. Wang, N. Gao, and Z. Huang, 2019. Structural basis of assembly of the human T cell receptor–CD3 complex. Nature 573:546–552.

39. Metropolis, N., A. W. Rosenbluth, M. N. Rosenbluth, A. H. Teller, and E. Teller, 1953. Equation of state calculations by fast computing machines. J Chem Phys 21:1087–1092.

40. Schroeder, D. V., 1999. An Introduction to Thermal Physics.

41. Press, W. H., S. a. Teukolsky, W. T. Vetterling, and B. P. Flannery, 2007. Numerical Recipes 3rd Edition: The Art of Scientific Computing.

42. Keul, N. D., K. Oruganty, E. T. Schaper Bergman, N. R. Beattie, W. E. McDonald, R. Kadirvelraj, M. L. Gross, R. S. Phillips, S. C. Harvey, and Z. A. Wood, 2018. The entropic force generated by intrinsically disordered segments tunes protein function.

43. Lopez, C. A., A. Sethi, B. Goldstein, B. S. Wilson, S. Gnanakaran, and C. A. Lo, 2015. Membrane-Mediated Regulation of the Intrinsically Disordered CD3E Cytoplasmic Tail of the TCR. Biophys J 108:2481–2491.

44. Ulmschneider, M. B., M. S. Sansom, and A. Di Nola, 2005. Properties of integral membrane protein structures: derivation of an implicit membrane potential. Proteins: Structure, Function, and Bioinformatics 59:252–265.

45. Portz, B., F. Lu, E. B. Gibbs, J. E. Mayfield, M. Rachel Mehaffey, Y. J. Zhang, J. S. Brodbelt, S. A. Showalter, and D. S. Gilmour, 2017. Structural heterogeneity in the intrinsically disordered RNA polymerase II C-terminal domain. Nat Comm 8:1–12.

46. Shi, X., Y. Bi, W. Yang, X. Guo, Y. Jiang, C. Wan, L. Li, Y. Bai, J. Guo, Y. Wang, X. Chen, B. Wu, H. Sun, W. Liu, J. Wang, and C. Xu, 2013. Ca2+ regulates T-cell receptor activation by modulating the charge property of lipids. Nature 493:111–115.

47. Ma, Y., Y. Yamamoto, P. R. Nicovich, J. Goyette, J. Rossy, J. J. Gooding, and K. Gaus, 2017. A FRET sensor enables quantitative measurements of membrane charges in live cells. Nat Biotech 35:363–370.

48. Ma, Y., K. Poole, J. Goyette, and K. Gaus, 2017. Introducing membrane charge and membrane potential to T cell signaling.

49. Wu, W., C. Yan, X. Shi, L. Li, W. Liu, and C. Xu, 2015. Lipid in T-cell receptor transmembrane signaling. Progress in Biophysics and Molecular Biology 118:130–138.

50. James, J. R., 2018. Tuning ITAM multiplicity on T cell receptors can control potency and selectivity to ligand density. Science Signaling 11.

51. Klammt, C., L. Novotná, D. T. Li, M. Wolf, A. Blount, K. Zhang, J. R. Fitchett, and B. F. Lillemeier, 2015. T cell receptor dwell times control the kinase activity of Zap70. Nat Immunol 16:961–9.

52. Liu, X., L. Bardwell, and Q. Nie, 2010. A combination of multisite phosphorylation and substrate sequestration produces switchlike responses. Biophys J 98:1396–1407.

53. Salazar, C., and T. Höfer, 2009. Multisite protein phosphorylation From molecular mechanisms to kinetic models. FEBS J 276:3177–3198.

54. Mukhopadhyay, H., S. P. Cordoba, P. K. Maini, P. A. van der Merwe, and O. Dushek, 2013. Systems Model of T Cell Receptor Proximal Signaling Reveals Emergent Ultrasensitivity. PLoS Comp Biol 9.

55. Suwanmajo, T., and J. Krishnan, 2015. Mixed mechanisms of multi-site phosphorylation. J R Soc Interface 12.

56. Wang, L., Q. Nie, and G. Enciso, 2010. Nonessential sites improve phosphorylation switch. Biophys J 99:L41–L43.

57. Rohrs, J. A., P. Wang, and S. D. Finley, 2019. Understanding the Dynamics of T-Cell Activation in Health and Disease Through the Lens of Computational Modeling. JCO Clinical Cancer Informatics 1–8.

58. Rangarajan, S., Y. He, Y. Chen, M. C. Kerzic, B. Ma, R. Gowthaman, B. G. Pierce, R. Nussinov, R. A. Mariuzza, and J. Orban, 2018. Peptide–MHC (pMHC) binding to a human antiviral T cell receptor induces long-range allosteric communication between pMHC- and CD3-binding sites. J Biol Chem 293:15991–16005.

59. Reboul, C. F., G. R. Meyer, B. T. Porebski, N. A. Borg, and A. M. Buckle, 2012. Epitope flexibility and dynamic footprint revealed by molecular dynamics of a pMHC-TCR complex. PLoS Comp Biol 8.

60. Huang, J., S. Rauscher, G. Nawrocki, T. Ran, M. Feig, B. L. De Groot, H. Grubmüller, and A. D. MacKerell, 2016. CHARMM36m: An improved force field for folded and intrinsically disordered proteins. Nat Methods 14:71–73.

61. Stanley, N., S. Esteban-Martín, and G. De Fabritiis, 2015. Progress in studying intrinsically disordered proteins with atomistic simulations. Progress in Biophysics and Molecular Biology 119:47–52.

62. Cortajarena, A. L., G. Lois, E. Sherman, C. S. O’Hern, L. Regan, and G. Haran, 2008. Non-random-coil Behavior as a Consequence of Extensive PPII Structure in the Denatured State. J Mol Biol 382:203–212.

63. Michalski, P. J., and L. M. Loew, 2016. SpringSaLaD: A Spatial, Particle-Based Biochemical Simulation Platform with Excluded Volume. Biophys J 110:523–529.

64. Hoffmann, M., C. Fröhner, and F. Noé, 2019. ReaDDy 2: Fast and flexible software framework for interacting-particle reaction dynamics. PLoS Comp Biol 15:e1006830.

65. Dyla, M., and M. Kjaergaard, 2020. Intrinsically disordered linkers control tethered kinases via effective concentration 117.

66. Neumeister Kersh, E., A. S. Shaw, and P. M. Allen, 1998. Fidelity of T cell activation through multistep T cell receptor ζ phosphorylation. Science 281:572–578.

67. Van Oers, N. S., B. Tohlen, B. Malissen, C. R. Moomaw, S. Afendis, and C. A. Slaughter, 2000. The 21- and 23-kD forms of TCRζ are generated by specific ITAM phosphorylations. Nat Immunol 1:322–328.

68. Germain, R. N., 1997. T-cell signaling: The importance of receptor clustering.

69. Sigalov, A. B., D. A. Aivazian, V. N. Uversky, and L. J. Stern, 2006. Lipid-binding activity of intrinsically unstructured cytoplasmic domains of multichain immune recognition receptor signaling subunits. Biochemistry 45:15731–15739.

70. Gagnon, E., D. a. Schubert, S. Gordo, H. H. Chu, and K. W. Wucherpfennig, 2012. Local changes in lipid environment of TCR microclusters regulate membrane binding by the CD3E cytoplasmic domain. J Exp Med 209:2423–39.

71. Koshland, D. E., and K. Hamadani, 2002. Proteomics and models for enzyme cooperativity. J Biol Chem 277:46841–46844.

