## Supplemental Text and Figures for "Intrinsic disorder in the T cell receptor creates cooperativity and controls ZAP70 binding"

#### List of Supporting Figures

S1 Local stiffening: Transition Rates . . . 4
S2 Local stiffening: Emergent sequential binding . . . 5
S3 Local stiffening: Unweighted average binding rates for short-range stiffening . . . 6
S4 Local stiffening: Binding rates for long-range stiffening . . . 7
S5 Local stiffening: Probability of second-to-last event . . . 8
S6 Local stiffening: Dose response curves and alternative Hill coefficient definitions . . . 9
S7 Membrane affinity: Zeta distributions . . . 10
S8 Membrane association: $\zeta$ binding rates and dose response curves . . . 11
S9 Membrane affinity: Dose response curves for $\epsilon$ alternative Hill coefficient definitions . . . 12
S10 Simultaneous binding: Filament sweep binding rates . . . 13
S11 Simultaneous binding: Filament sweep binding rates histograms . . . 14
S12 Integrative model full results . . . 15

#### List of Supporting Text

**1 Summary of interactions** **2**

**2 Local stiffening** **2**

**3 Ultrasensitivity and alternate definitions of Hill coefficients** **2**

**4 Multimodality in binding rates** **3**

**5 List of Supporting Movies** **3**

### 1 Summary of interactions

To summarize interactions between physical elements: Ligands are treated as hard spheres, so there is zero repulsion until they approach the chain, or each other, until within 1 radius, after which there is infinite repulsion. Bound ligands are constrained to move with the chain, equivalent to an infinitely-strong bond. Interactions with the membrane are modeled using the general forms in Main Text Eqs. 5-6, where the parameters $E_{P0}$ and $E_{B0}$ are estimated by constraining the model to satisfy two experimental observations (tyrosine phosphorylation and basic residue mutation). Within-chain interactions are omitted, i.e., it is a non-interacting FJC.

### 2 Local stiffening

As described in Main Text Fig. 4, we explore the consequences of assuming that a phosphorylation event locally stiffens the region around the tyrosine. Here, we explore stiffening ranges beyond 11 amino acids in Fig. S4. For maximal local stiffening, i.e. all residues becoming stiff upon first phosphorylation, the binding rate increases to 1.0 in units relative to the free-space (unoccluded) rate, since the kinase is uninhibited and always able to access the binding sites.

At intermediate values of local stiffening, the average binding rate does not consistently increase but oscillates (Fig. S4A). This phenomenon can be understood by considering the distribution of tyrosines along $\zeta$. The six tyrosines are grouped in pairs, constituting ITAMs. The tyrosines in each pair are significantly closer to each other than to the tyrosines of the next ITAM, leading to an evens-odds effect in which even-numbered tyrosines are fast compared to odd-numbered tyrosines.

The Main Text figures weight the phosphorylation rates by their likelihood. Without weighting by most likely phosphorylation states, the average binding rate smoothly increases with number of phosphorylations, as shown in Fig. S3.

### 3 Ultrasensitivity and alternate definitions of Hill coefficients

Here we investigate two complementary defintions of ultrasensitivity.Both definitions extract a Hill coefficient from the dose-respone curves. We refer to them here as switch-like behavior and threshold behavior. The switch-like behavior of a dose-response curve describes how quickly it increases from a less-phosphorylated state to a more-phosphorylated state and can be used as a measure of cooperativity. Mathematically, the switch-like behavior is the maximum logarithmic slope of each curve. Specifically, for a kinase-phosphatase ratio $r$, if the fraction of tyrosines phosphoryated is $p(r)$, then

$$H_{\text{switch}} = \max\left(\frac{d}{dr}\left(\log\left(\frac{p(r)}{1-p(r)}\right)\right)\right). \tag{1}$$

Alternatively, the threshold behavior of the curve describes how high of a dose is required before a response occurs. We calculate this value as

$$H_{\text{threshold}} = \log_{10}(81)/\log_{10}(\text{EC90}/\text{EC10}). \tag{2}$$

We calculate the switch-like and threshold behavior of the dose-response curves under constant and steric dephosphorylation in Fig. S6.

For constant dephosphorylation, the switch-like behavior increases with the number of amino acids stiffened per phosphorylation event. When 1/12 chain is stiffened per event, the switch hill coefficient is 1.8, suggesting moderate switch-like behavior, while the threshold hill coefficient is less than 1.2, suggesting a poor threshold (Fig. S6C). For steric dephosphorylation, the same increasing trend persists, but at 1/12 chain local stiffening, the switch hill coefficient is only 1.3 and the threshold hill coefficient is about 0.8 (Fig. S6D). Local stiffening therefore creates cooperativity, even under reversible phosphorylation, but makes a poor threshold.

In Fig. S9 we show the Hill coefficients for the $\epsilon$ membrane affinity model presented in the Main Text, but with the threshold definition of Hill coefficient in Eq. 2.

### 4 Multimodality in binding rates

We simulate simultaneous ligand binding to 10 sites distributed among 1, 5, and 10 chains. Specifically, we consider the rate of binding a sixth ligand to the complex. Under this construction, there are 1260 ways (= 5 $\times$ 10 choose 5) a sixth ligand may bind to the complex. Histograms of these 1260 binding rates for each complex construction show that multimodality can arise in some cases. When there are 5 filaments, with 2 binding sites each, we see bimodality in the binding rates when the space between binding sites is 8 amino acids (Fig. S11). We find weak tri-modality when the binding site spacing is longer (20 amino acids).

To explore how this multi-modality arises, we investigated characteristics of the binding sites. We split the binding sites into two categories: binding sites on a filament with no ligands bound and binding sites on a filament that already has a ligand bound. When we plot the histograms of the binding rates by category, we find that the bimodal distribution can be explained. Binding a second ligand to a filament is harder than binding the first ligand, since the second filament will experience steric occlusion from the other ligand in addition to the filament itself. This results in a lower binding rate compared to ligands binding to empty filaments (Fig. S11). Steric occlusion from ligands on the same filament is increased when the space between the binding sites is shorter. This explains the high separation between the two modes of binding rates for the shorter binding site spacing compared to the longer binding site spacing.

### 5 List of Supporting Movies

- Supporting Movie SM1: Example of simulated TCR with two bound ligands. The chains of the TCR are shown simulated as freely-jointed chains ($\zeta$ - green, $\epsilon$ - red, $\delta$ - blue, $\gamma$ - purple) with ITAM binding sites shown as black squares. Chains are anchored (black x) in a narrow base configuration, $\sim 1.5\,$nm apart and not allowed to pass below the membrane (yellow). Two bound ligands (opaque orange spheres) are shown attached to TCR. A third ligand attempts to bind a specific binding site. When binding region is occluded in a configuration, ligand cannot bind (transparent orange sphere). When polymers, bound ligands, and membrane are all outside of binding region, the ligand may bind (not shown). Radius of ligands is 2.1 nm.

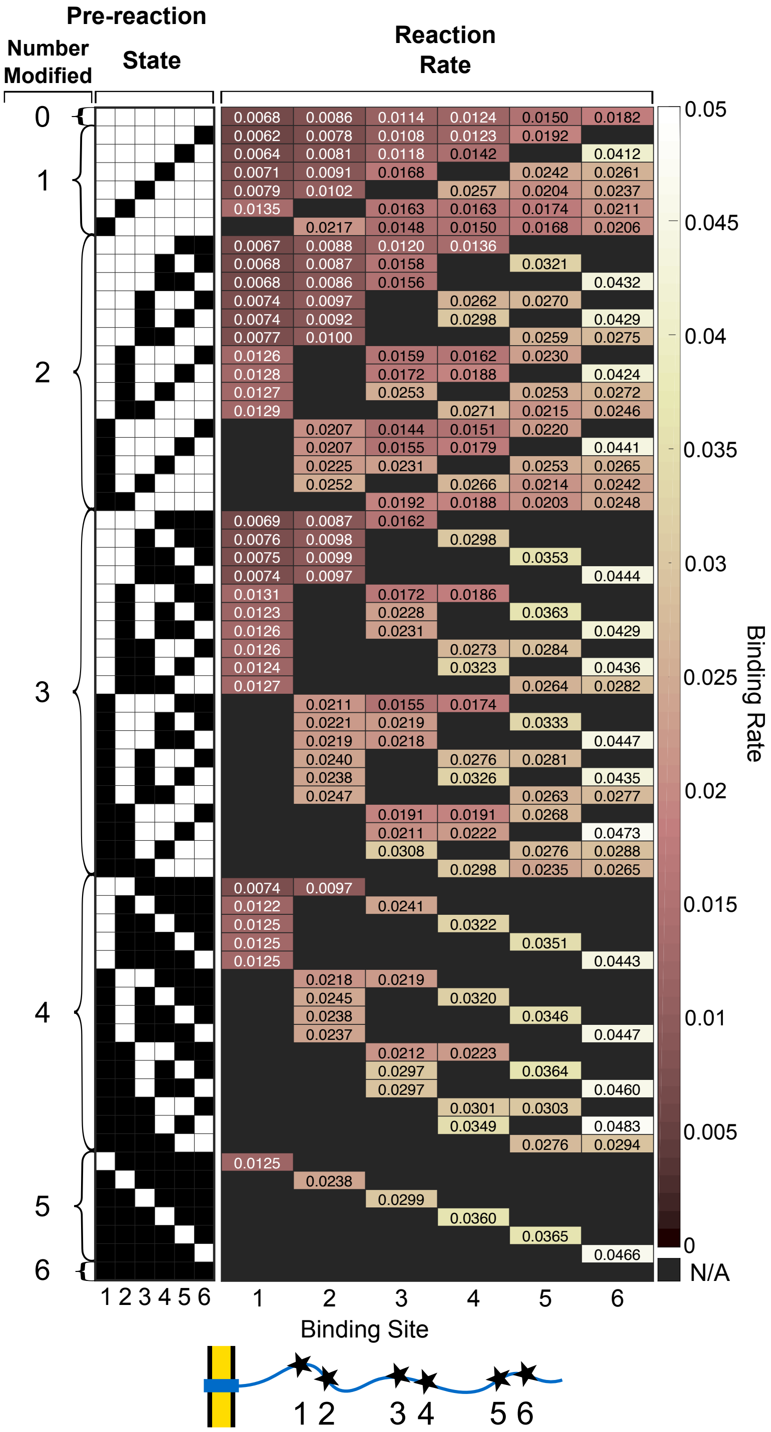


Fig. S1: **Disorder leads to accessibility differences, which implies binding rate differences between phosphorylation sites.** Reaction rates of a kinase binding to the six binding sites of membrane bound $\zeta$ in a given phosphorylation state. In this case, results are shown assuming each phosphorylation event locally stiffens 11 amino acids. Dark red: low binding rate; white: high binding rate; black: phosphorylated site. Left two columns show the pre-reaction number modified and phosphorylation state before the next binding event (white: unphosphorylated site; black: phosphorylated site).

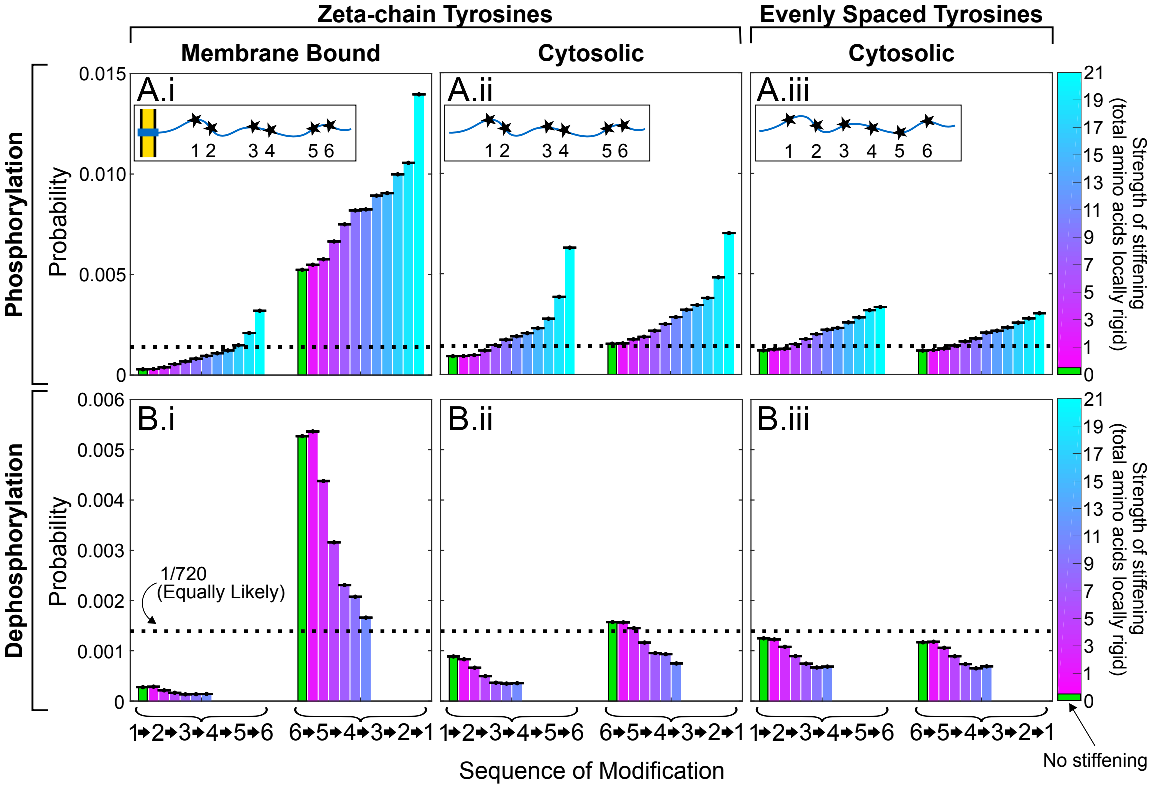


Fig. S2: **Binding rate differences imply emergent preference of (de)phosphorylation sequence.** Probability of (A) phosphorylating or (B) dephosphorylating membrane proximal-to-distal (123456) compared to membrane distal-to-proximal (654321). Colors represent simulations of different ranges of local stiffening/unstiffening per binding event. No stiffening (0 stiffening) shown in green. Each of these is shown for: (i) with a membrane, assuming sites spaced like tyrosines of $\zeta$, (ii) without a membrane, assuming sites spaced like tyrosines of $\zeta$, and (iii) without a membrane, assuming sites spaced evenly along the length of $\zeta$. The black dotted lines indicate the probability if all events were equally likely ($1/6!{=}1/720$). Error bars (A,B) represent standard error of the mean, treating each Gillespie run as an individual Bernoulli trial. Phosphorylation and dephosphorylation are more likely to occur membrane distal-to-proximal compared to proximal-to-distal for both the membrane bound and cytosolic $\zeta$ chain. For cytosolic $\zeta$ with evenly spaced tyrosines, both (de)phosphorylation sequences are approximately equally probable.

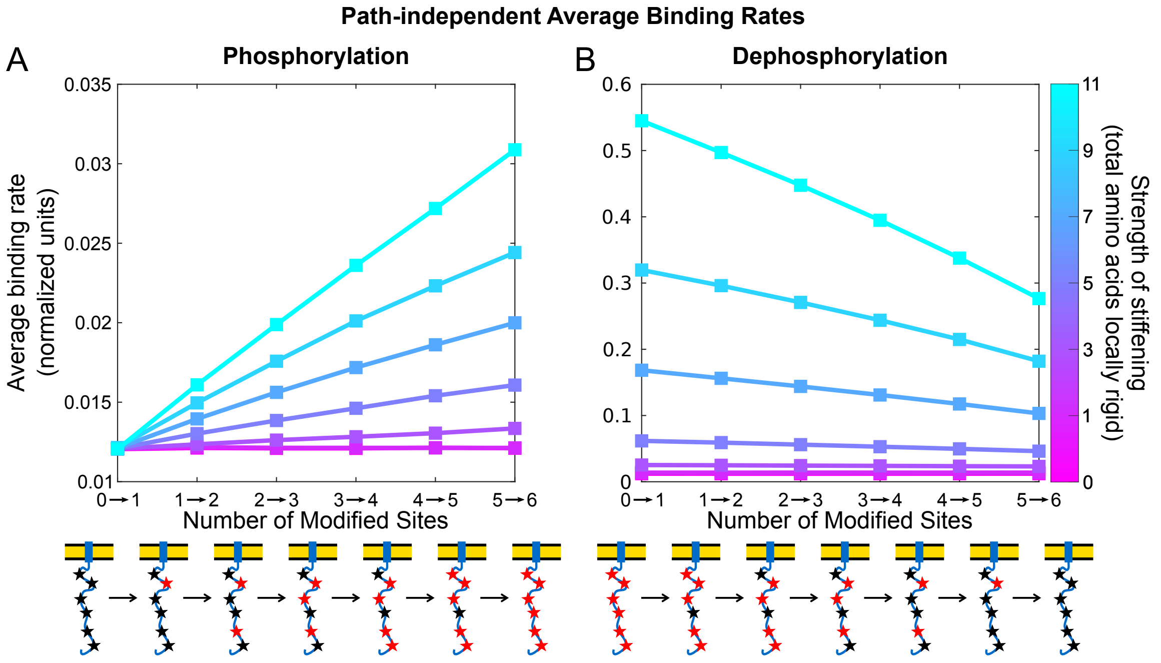


Fig. S3: **Local stiffening: Unweighted average binding rates for short-range stiffening** (A) Sequence-independent average binding rates of a kinase binding to $\zeta$ at different phosphorylation states for varying range of local stiffening (pink: no residues are stiffened; blue: 11 residues are stiffened, 5 on each side of site). The kinase average binding rate increases with each phosphorylation and also with local stiffening range. (B) Sequence-independent average binding rates of a phosphatase binding to $\zeta$ at different dephosphorylation states for varying ranges of local un-stiffening per dephosphorylation event. When local stiffening occurs, the phosphatase average binding rate decreases with each phosphorylation and also with local stiffening level. For both (A) and (B), schematic below axis shows example configuration for each phosphorylation state. Unphosphorylated residues represented by black stars, phosphorylated residues are red stars. Kinase and phosphatase radius is 2.1nm. Rates are normalized to the free-space binding rate.

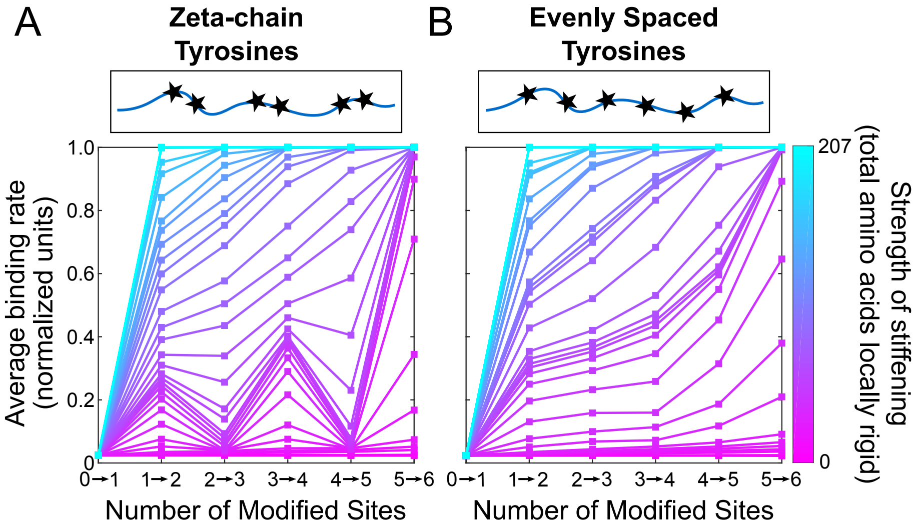


Fig. S4: **Local stiffening: Binding rates for long-range stiffening.** Sequence-dependent average binding rates of a kinase binding to cytosolic $\zeta$ (A) and cytosolic $\zeta$ with evenly spaced tyrosines (B) at different phosphorylation states for varying range of local stiffening (pink: no residues are stiffened; blue: all residues are stiffened after first phosphorylation). For small and large magnitudes of local stiffening, the average binding rate increases with each phosphorylation. For intermediate magnitudes of local stiffening, cytosolic $\zeta$ exhibits oscillations in the average binding rate with an overall increase (A) while cytosolic $\zeta$ with evenly spaced tyrosines increases without oscillations (B). In both cases at maximal local stiffening per phosphorylation, the entire domain is stiff leading to uninhibited kinase binding. Schematic below axis shows example configuration for each phosphorylation state. Unphosphorylated residues represented by black stars, phosphorylated residues are red stars. Kinase and phosphatase radius is 2.1nm. Rates are normalized to the free-space binding rate.

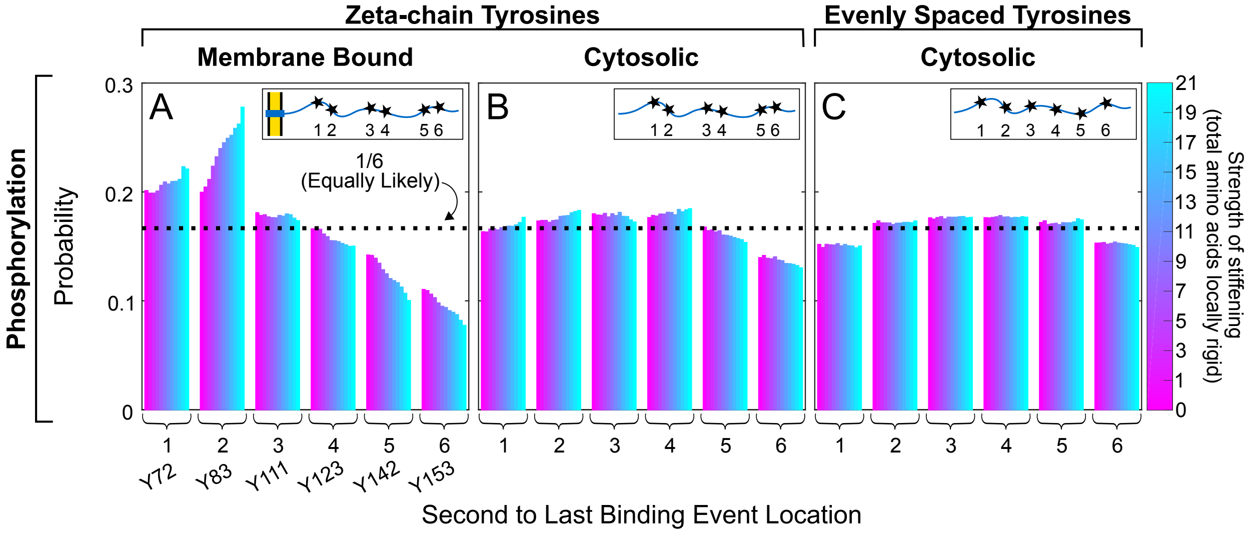


Fig. S5: **Local stiffening: Probability of second-to-last event.** Probability of the second-to-last binding event occurring at a specific site. Colors represent simulations of different ranges of local stiffening/unstiffening per binding event. Shown for: (A) with a membrane, assuming sites spaced like tyrosines of $\zeta$, (B) without a membrane, assuming sites spaced like tyrosines of $\zeta$, and (C) without a membrane, assuming sites spaced evenly along the length of $\zeta$. The black dotted lines indicate the probability if all events were equally likely (1/6). As reported in Main Text Fig. 3, there is a decrease in the average binding rate between the fourth and fifth binding event. To understand this effect, we examine the probability of the identity of the second-to-last binding event For membrane-bound $\zeta$-chains, the fifth binding event is most likely to occur at the second tyrosine from from the membrane, Y83. Therefore, the average binding rate is most heavily influenced by the binding rate at the Y83. The dip in binding rate seen there marks a competition between disordered-to-ordered transitions, which tends to make all sites more accessible, versus the increasing difficulty of phosphorylating sites that are highly occluded. Although occlusion by the chain is reduced by disordered-to-ordered transitions, it is not enough to overcome the high membrane occlusion that Y83 experiences. Therefore, since the fifth binding event is dominated by the low binding rate that Y83, the average rate decreases.

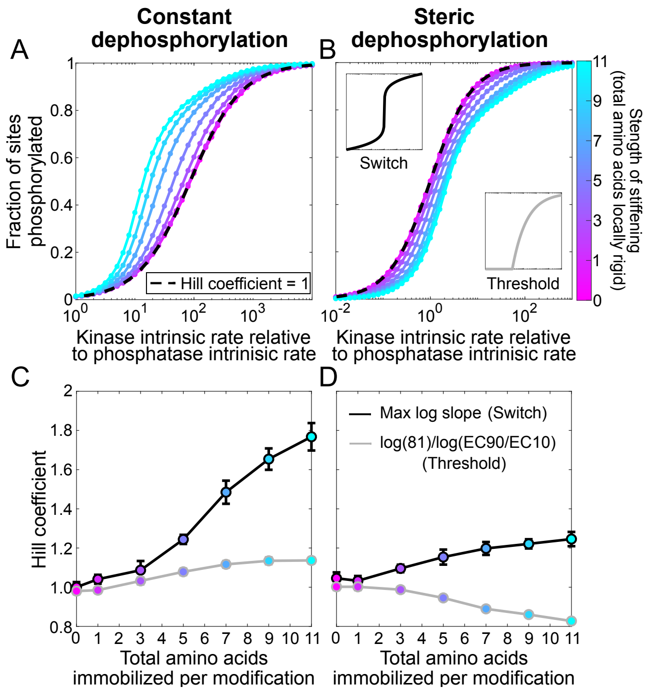


Fig. S6: **Local stiffening: Dose response curves and alternative Hill coefficient definitions.** (A,B) Fraction of sites phosphorylated as a function of the kinase-to-phosphatase activity ratio (measured by the ratio of free-space rates) for different ranges of local stiffening (colorbar), assuming a phosphatase with (A) negligible size or (B) radius of 2.1 nm, equal to the kinase. Black dashed line indicates a linear dose-response, i.e., with Hill coefficient 1. (C,D) Hill coefficients for different ranges of local stiffening per phosphorylation event, assuming a phosphatase of (C) negligible size or (D) radius of 2.1 nm. Hill coefficients calculated from maximum log-log slope (black line) or log(81)/log(EC90/EC10) (gray line) of the dose response curves. Error bars for max log slope indicate root-mean-square error from a cubic polynomial fit to slope. Error bars for log(81)/log(EC90/EC10) indicate standard deviation of hill coefficients from bootstrap sampling from dose-response curve.

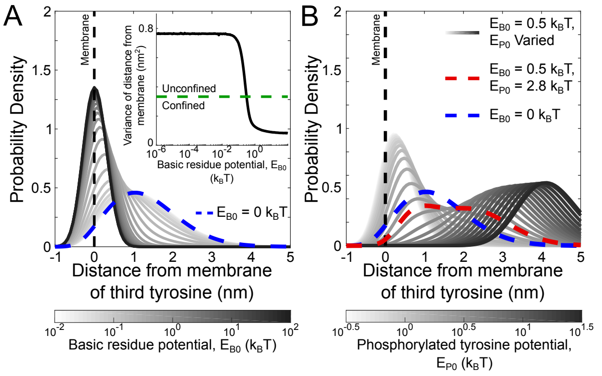


Fig. S7: **Simplified membrane interaction model sets constraints on the strength of basic residue attraction and phosphorylation-driven repulsion from membrane.** (A) Probability density of the distance from the membrane of the 3rd tyrosine of $\zeta$, assuming no phosphorylated tyrosines, for varying strengths of the basic residue potential, $E_{\mathrm{B0}}$ (white: low; black: high). The tyrosine moves close to the membrane as $E_{\mathrm{B0}}$ is increased. Probability density when there is no basic residue potential ($E_{\mathrm{B0}} = 0\ k_{\mathrm{B}}T$) is shown as blue dotted line. (Inset) Variance of probability density of 3rd tyrosine over range of basic residue potential strengths. Green dashed line shows the characteristic $E_{\mathrm{B0}} = 0.5 k_{\mathrm{B}}T$ required to confine the tyrosine to the membrane, defined in the text. (B) Probability density of the location of 3rd tyrosine assuming all tyrosines are phosphorylated, for $E_{\mathrm{B0}} = 0.5\ k_{\mathrm{B}}T$ (the value required to confine the tyrosine to the membrane assuming no tyrosines are phosphorylated), and varying phosphorylated tyrosine potential strength, $E_{\mathrm{P0}}$ (low - white, high - black). Probability density when $E_{\mathrm{P0}} = 2.8\ k_{\mathrm{B}}T$ is shown as red dashed line, reflecting the $E_{\mathrm{P0}}$ value needed to approximately return to the distribution when $E_{\mathrm{B0}} = 0\ k_{\mathrm{B}}T$ (blue dashed line).

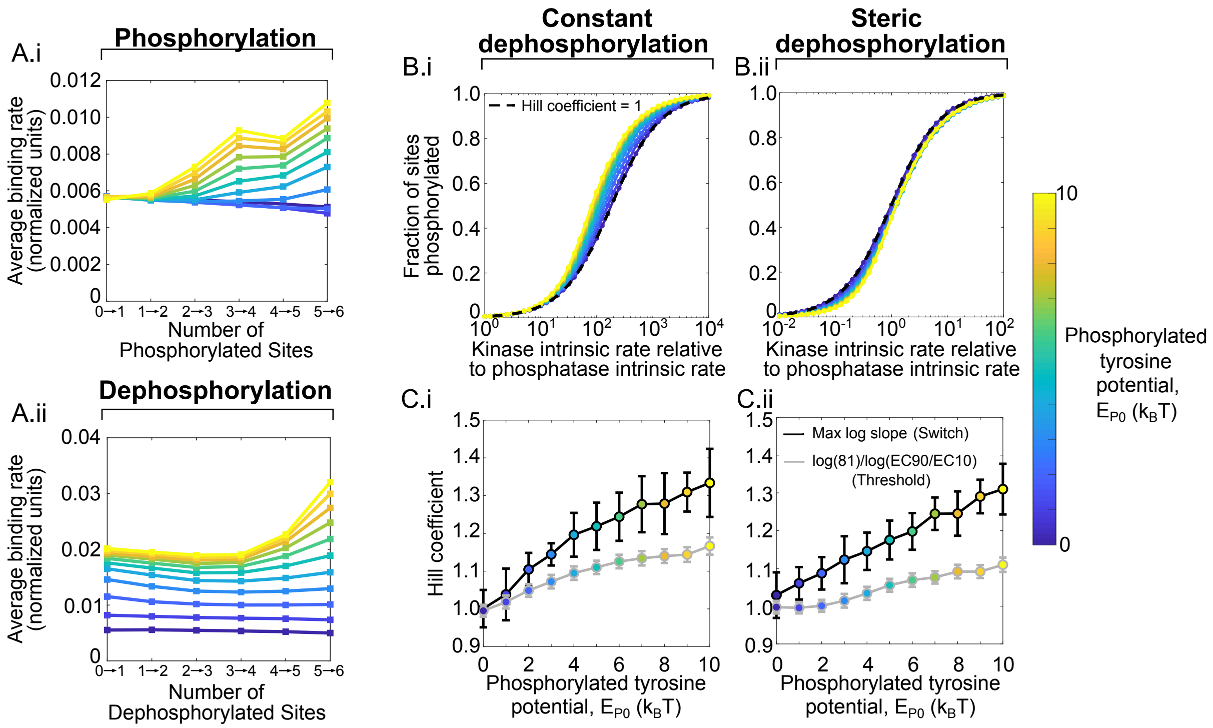


Fig. S8: **Phosphorylation-driven modulation of membrane association of $\zeta$ leads to cooperativity and sequential binding.** (A) Sequence-dependent average binding rates of (i) kinase and (ii) phosphatase binding to $\zeta$ at different (de)phosphorylation states for varying strengths of phosphorylated tyrosine potential ($E_{\mathrm{P0}}$) (blue: weak; yellow: strong). Both kinase and phosphatase binding rates increase with each phosphorylation and also with increasing $E_{\mathrm{P0}}$. (B) Fraction of sites phosphorylated over kinase intrinsic rate compared to phosphatase intrinsic rate for varying strengths of phosphorylated tyrosine potential ($E_{\mathrm{P0}}$), assuming a phosphatase with (i) negligible size (constant dephosphorylation) or (ii) 2.1 nm radius, equivalent to the kinase (steric dephosphorylation). Black dashed line indicates linear dose response, i.e., Hill coefficient 1. (C) Hill coefficients for varying strengths of phosphorylated tyrosine potential ($E_{\mathrm{P0}}$), assuming a phosphatase of (i) negligible size or (ii) radius of 2.1 nm, equal to the kinase. Hill coefficients calculated from maximum log-log slope (black line) or log(81)/log(EC90/EC10) (gray line) of the dose response curves. Error bars for max log slope indicate root-mean-square error from a cubic polynomial fit to slope. Error bars for log(81)/log(EC90/EC10) indicate standard deviation of hill coefficients from bootstrap sampling from dose-response curve. In both cases, the Hill coefficient increases with the strength of phosphorylated tyrosine potential, even when dephosphorylation is assumed to be sterically hindered. For (A)-(C), both kinase and phosphatase are assumed to have radius 2.1 nm. Rates are normalized to the free-space binding rate.

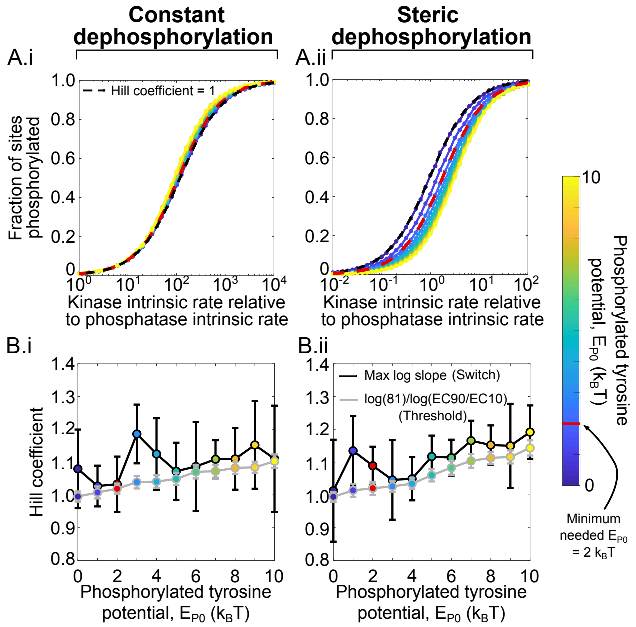


Fig. S9: **Membrane affinity: Dose response curves for $\epsilon$ alternative Hill coefficient definitions.** (A) Fraction of sites phosphorylated over kinase intrinsic rate compared to phosphatase intrinsic rate for varying strengths of phosphorylated tyrosine potential ($E_{\mathrm{P0}}$), assuming a phosphatase with (i) negligible size (constant dephosphorylation) or (ii) 2.1 nm radius, equivalent to the kinase (steric dephosphorylation). Black dashed line indicates linear dose response, i.e., Hill coefficient 1. (B) Hill coefficients for varying strengths of phosphorylated tyrosine potential ($E_{\mathrm{P0}}$), assuming a phosphatase of (i) negligible size or (ii) radius of 2.1 nm, equal to the kinase. Hill coefficients calculated from maximum log-log slope (black line) or log(81)/log(EC90/EC10) (gray line) of the dose response curves. Error bars for max log slope indicate root-mean-square error from a cubic polynomial fit to slope. Error bars for log(81)/log(EC90/EC10) indicate standard deviation of hill coefficients from bootstrap sampling from dose-response curve. In both cases, the Hill coefficient increases with the strength of phosphorylated tyrosine potential, even when dephosphorylation is assumed to be sterically hindered (B). Rates are normalized to the free-space binding rate.

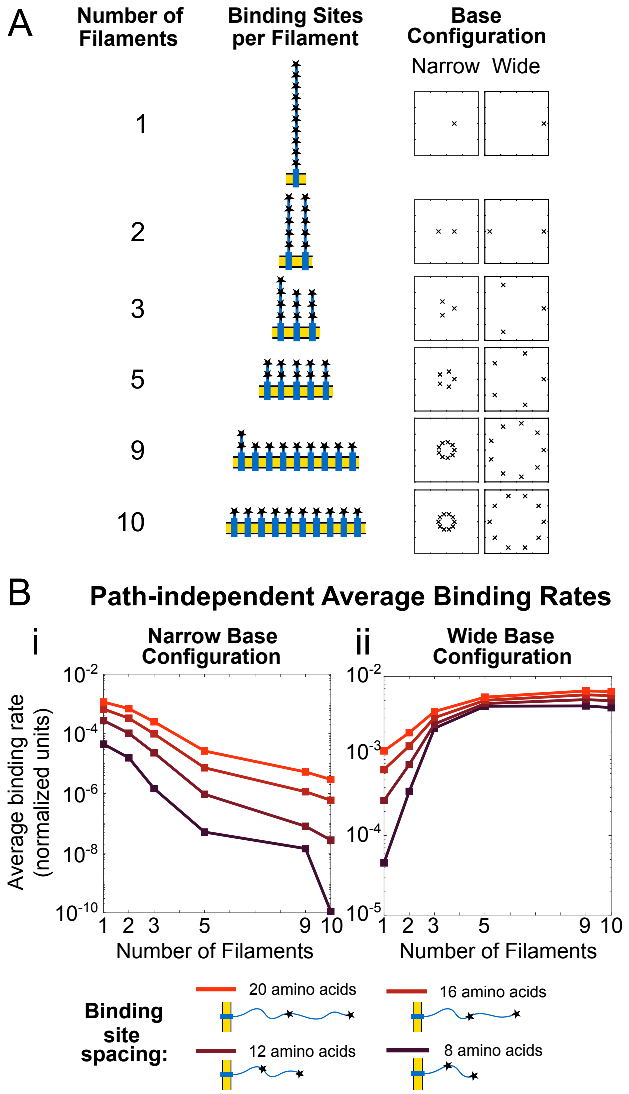


Fig. S10: **Optimal distribution of ten binding sites on multiple chains is dependent on membrane spacing of subunits.** (A) Schematic for distribution of 10 binding sites on multiple chains. For each, filaments are distributed evenly on a (1) narrow circle of radius 1.5 nm, and (2) wide circle of radius 5 nm. (B) Average binding rates of sixth binding event to constructed domain against number of filaments in domain (color bar; dark red: short spacing between binding sites; bright red: long spacing) and subunit configuration (Bi) 1.5nm radius, (Bii) 5nm radius. Simulations sweep over binding site spacing from 8 to 20 amino acids, indicated by color. Ligand radius is 2.7 nm.

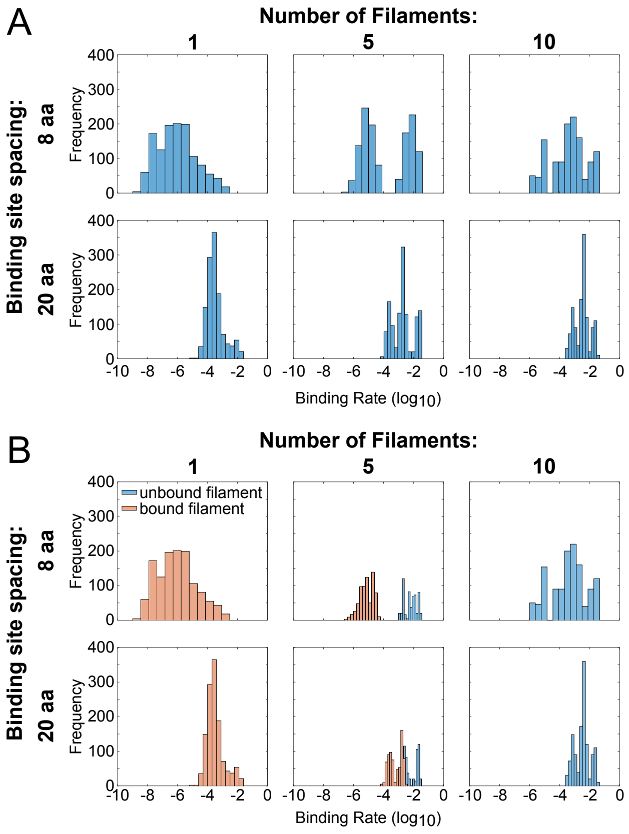


Fig. S11: **Bimodality of binding rates can be explained by ligands bound to target filament.** (A) Histograms of binding rates of sixth binding event to constructed domains in wide base configuration. (B) Histograms of binding rates from (A) colored by binding to filaments with no ligands bound (blue) and filaments with ligands bound (orange). Ligand radius is 2.7 nm.

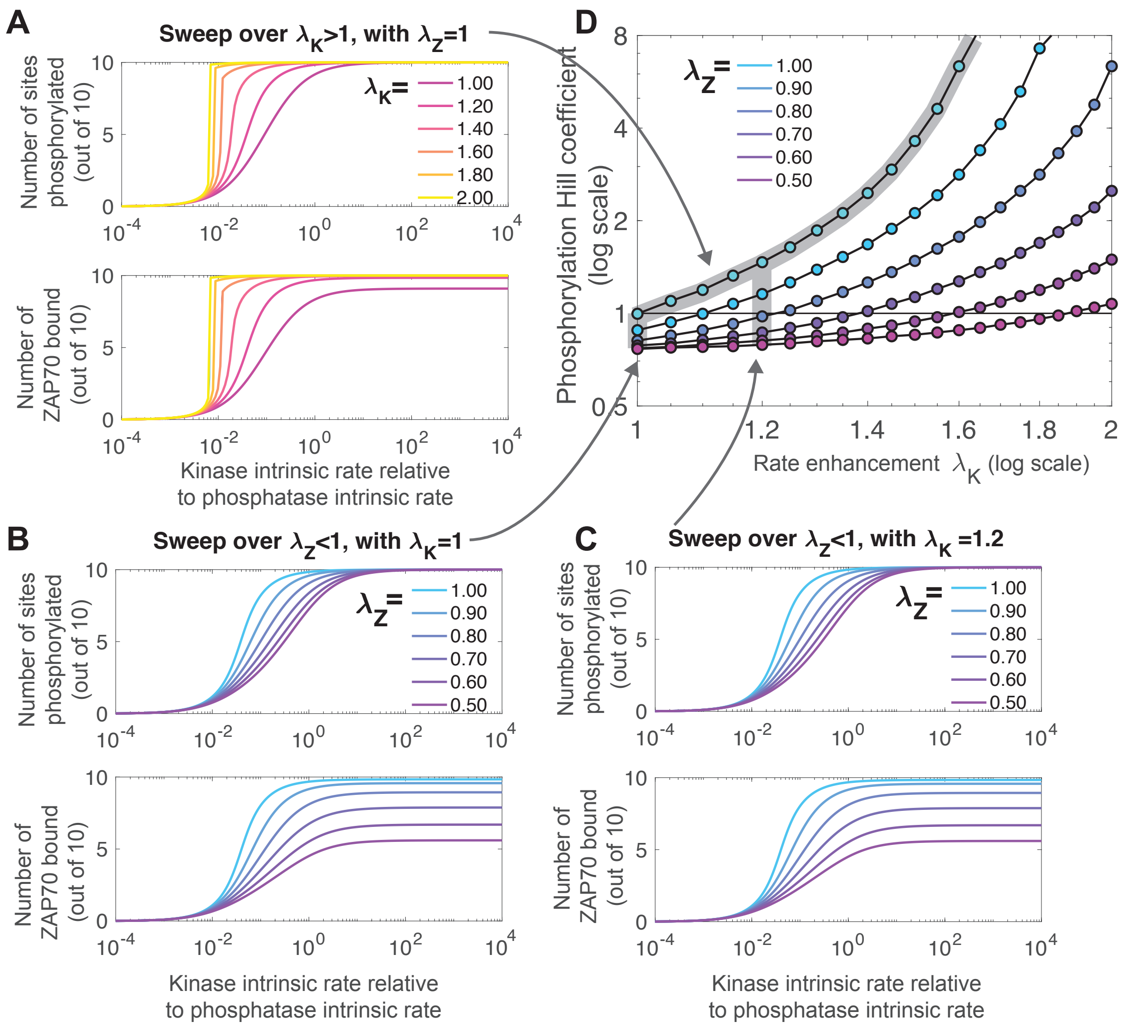


Fig. S12: **Full results of integrative model shows counteracting effects of rate enhancement and rate reduction.** (A) Rate enhancement of kinase phosphorylating TCR $\lambda_K > 1$ leads to more switch-like dose response of both amount of phosphorylation (top) and amount of ZAP70 bound (bottom). (B) Rate decrease of ZAP70 binding $\lambda_Z < 1$ leads to shallow dose-response for both amount of phosphorylation and amount of ZAP70 bound. In addition, for ZAP70-bound, the rate decrease also leads to a lower saturating value, i.e., $EC_{\max}$. (C) When both rate enhancement ($\lambda_K = 1.2$) and rate decrease ($\lambda_Z < 1$, different values explored) are present, the switch-like response is abrogated. (D) Ultrasensitivity quantified using the log-effective-concentration definition Eq. 2 for a range of $\lambda_K \geq 1$ and a range of $\lambda_Z \leq 1$.
